# Pantothenate Kinase 4 controls efficient skeletal muscle energy substrate metabolism via acetyl-CoA

**DOI:** 10.1101/2023.08.14.551603

**Authors:** Adriana Miranda-Cervantes, Andreas M. Fritzen, Steffen H. Raun, Ondřej Hodek, Lisbeth L. V. Møller, Kornelia Johann, Luisa Deisen, Paul Gregorevic, Anders Gudiksen, Anna Artati, Jerzy Adamski, Nicoline R. Andersen, Peter Schjerling, Alberto Cebrian-Serrano, Markus Jähnert, Pascal Gottmann, Ingo Burtscher, Heiko Lickert, Henriette Pilegaard, Annette Schürmann, Matthias H. Tschöp, Thomas Moritz, Timo D. Müller, Lykke Sylow, Bente Kiens, Erik A. Richter, Maximilian Kleinert

## Abstract

Metabolic inflexibility in skeletal muscle (SkM) is closely linked to metabolic diseases. Exercise improves metabolic flexibility, rendering it a valuable discovery tool of mechanisms promoting efficient metabolism of glucose and lipids. We herein discover pantothenate kinase 4 (PanK4) as a conserved exercise target with high abundance in SkM. We go on to show that murine muscle *Pank4* is dysregulated with high-fat diet feeding, and identify human *PANK4* variants that associate with glycemic control and body mass index traits, indicating important roles of PanK4 in glucose metabolism and growth. Consistent with the latter, germline deletion of PanK4 reduces circulating IGF-1 and stunts growth in mice. Deletion specifically in mouse SkM reveals that PanK4 facilitates fatty acid oxidation by acting as a regulator of SkM acetyl-CoA, a key node in metabolism of both glucose and lipids. Consequently, without PanK4, elevated SkM acetyl-CoA levels allosterically gridlock key enzymes required for efficient lipid and glucose utilization, and these SkM metabolic perturbations manifest in whole-body insulin resistance. As proof of principle, we show that an increase in muscle PanK4 lowers SkM acetyl-CoA and increases SkM glucose utilization. Our findings identify PanK4 as a novel regulator of SkM energy substrate metabolism, warranting inclusion in comprehensive strategies against metabolic disease.

## Introduction

Skeletal muscle (SkM) makes up approximately 30-40% of body mass^1^ and plays a crucial role in maintaining overall glucose balance in the body. In normal circumstances, SkM is responsible for the majority of glucose uptake stimulated by insulin^2^ and serves as the primary storage site for glucose in the form of glycogen^3^. Therefore, any metabolic conditions that impair the ability of SkM to utilize glucose efficiently pose a significant challenge to maintaining overall glucose homeostasis in the body.

Obesity is closely associated with insulin resistance of SkM, which is considered a primary risk factor for the development of type 2 diabetes (T2D)^4^. Obesity-induced insulin resistance in SkM involves complex mechanisms, including inflammation, mitochondrial dysfunction, endoplasmic reticulum stress, and lipotoxicity^5–7^. In obesity, adipocytes exceed their lipid storage capacity, releasing fatty acids and cytokines into the bloodstream. These harmful lipids accumulate in non-adipose tissues, causing local inflammation and contributing to insulin resistance development. In addition, it has been suggested that obesity induces metabolic inflexibility in SkM^8^. Normally, we experience alternating periods of feeding and fasting, necessitating metabolic flexibility to switch between using glucose and fatty acids as energy sources. The reciprocal relationship between glucose and fatty acid was first 60 years ago^9^ but the underpinning mechanism are not fully elucidated. With prolonged and frequent overeating an excess supply of glucose and fatty acids to SkM cells causes mitochondrial congestion^10^, derailing the coordinated utilization of glucose and fatty acids. This metabolic perturbation is thought to contribute to insulin resistance in SkM and challenges systemic glucose homeostasis.

Exercise has been found to increase glucose uptake in skeletal muscle in both healthy individuals and those with insulin resistance^11^, making it an effective intervention for reducing blood glucose levels in people with type 2 diabetes. Furthermore, exercise has been demonstrated to improve the metabolic flexibility of skeletal muscle^8^ and enhance insulin sensitivity^12^, thereby contributing to better overall glucose balance in the body. Thus, exercise-associated signaling mechanisms include potential therapeutic targets that regulate efficient utilization of glucose and lipids, some of which may still be unidentified. We show that pantothenate kinase 4 (PanK4) is a new exercise target abundant in both rodent and human skeletal muscle. Moreover, we demonstrate that PanK4 plays a crucial role in coordinating efficient utilization of glucose and lipids in SkM.

## Results

### Pantothenate kinase 4 (PanK4) is a new exercise target in skeletal muscle

Datasets on exercise- or contraction-induced changes in protein phosphorylation^13, 14^ indicate that pantothenate kinase 4 (PanK4) has a conserved phosphorylation site at Ser63 (p-PanK4^Ser63^) that is acutely increased by exercise/muscle contractions in rodent and human SkM (Fig 1a). PanK4 belongs to the family of pantothenate kinases (PanK1-4) that regulate the biosynthesis of coenzyme A (CoA), a pivotal coenzyme required in many cellular processes, including the synthesis, storage and uptake of fatty acids (FAs)^15^. However, it has recently emerged that PanK4 does not exhibit classical pantothenate kinase activity^16^, but instead may rather antagonize the production of coenzyme A by acting as a phosphatase^17^. This is consistent with the pantothenate kinase domain sequence found in PanK4 being significantly different from that of its family members, PanK1-3 (Fig S1a). Clearly, PanK4 is unique but its metabolic and physiological function are unknown.

**Figure 1.**
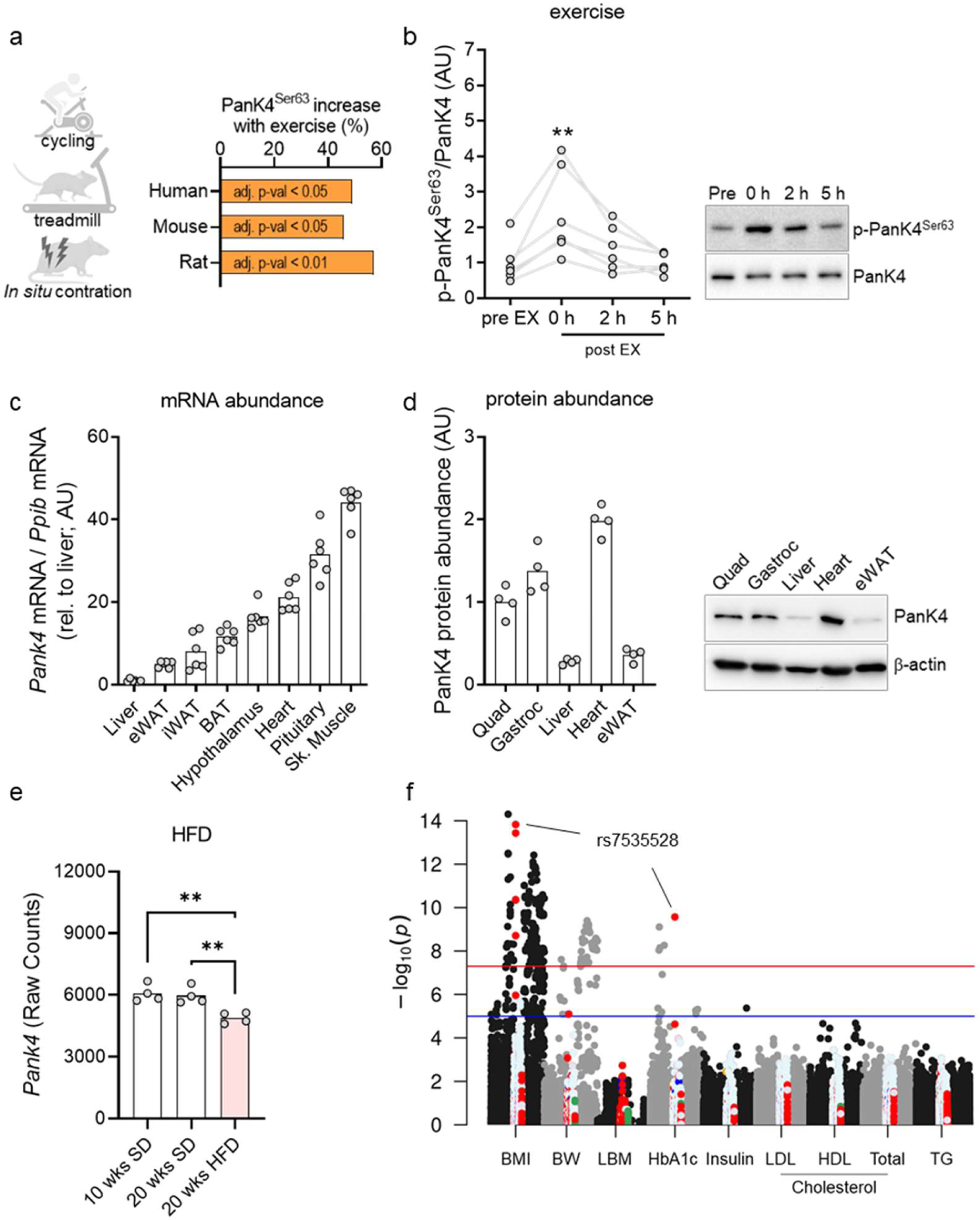
PanK4 is an exercise target abundant in skeletal muscle and associated with metabolic dysregulation. **a,** PanK4 Ser63 phosphorylation (p-PanK4^Ser63^) site and its regulation by exercise or muscle contraction in skeletal muscle from humans and rodents identified by phosphoproteomics^13, 14^. **b,** p-PanK4^Ser63^ in human vastus lateralis before, immediately post and at indicated time-points into recovery from vigorous cycling exercise. **c,d** *Pank4* mRNA and PanK4 protein abundance in indicated tissues**. e,** murine *Pank4* in skeletal muscle from mice fed a standard diet (SD) or a high-fat diet for indicated number of weeks. **f,** single-nucleotide polymorphisms (SNPs) 50kb up- and downstream of the human *PANK4* gene and their associations with indicated traits. Red circles are SNPs within *PANK4*. Red horizontal line indicates threshold for genome wide association. **p<0.01 vs Pre-EX or as indicated. Statistic: b, repeated measures one-way ANOVA; e, one-way ANOVA. Statistical analyses were conducted with log2-transformed data. Šidák post hoc testing was performed whenever respective ANOVA yielded significance.

To verify the phosphoproteome discovery, we generated and extensively validated a phospho-specific antibody against p-PanK4^Ser63^ (Fig S1b,c). Using this antibody, we show that after vigorous cycling exercise in humans (∼10 min at 77%-88% *W*_max_), p-PanK4^Ser63^ increased 2.5-fold in vastus lateralis muscle and returned to pre-exercise levels five hours later (Fig 1b). These data confirm that PanK4 is a novel exercise target in human skeletal muscle.

### PanK4 is abundant in striated muscle and its expression associates with conditions of metabolic dysregulation

We next sought to understand more about Pank4 expression. We detected high levels of *Pank4* mRNA in skeletal and cardiac muscle, hypothalamus and the pituitary gland of mice (Fig 1c). We also found that protein abundance of PanK4 in mice is high in skeletal muscle and heart compared to liver and adipose tissue (Fig 1d). Murine *Pank4* mRNA abundance was decreased in SkM from 20 weeks high-fat diet-induced mice with obesity (Fig 1e)^18^, which is known to be associated with severe insulin resistance^19^. Additionally, we identified human *PANK4* variants that are highly significantly linked to glycated hemoglobin (HbA_1c_) and body mass index (BMI) traits (Fig 1f). Collectively, these findings highlight a significant abundance of PanK4 in SkM and suggest the potential involvement of PanK4 in the regulation of growth and glucose metabolism.

### Germline deletion of PanK4 reduced circulating IGF-1 and stunted growth

To investigate the metabolic function of PanK4, we generated whole-body PanK4 knockout (PanK4 KO) mice by replacing the open reading frame of *Pank4* by a NLS-lacZ-2A-H2B-Venus cassette^20^. Both male and female PanK4 KO mice had lighter skeletal muscles based on wet weight mass (Fig S2c,d). Female PanK4 KO mice also had lower body weight, smaller hearts, livers, were overall shorter and had reduced levels of circulating insulin-like growth factor 1 (IGF-1) (Fig S2b,d,f). Male PanK4 KO mice had similar body weight and length as their WT counterparts, but their heart mass tended (p < 0.1) to be lower and circulating IGF-1 levels were reduced relative to WT mice as in female PanK4 KO mice (Fig S2a,c,e). These data indicate that PanK4 is critical for organismal development.

### PanK4 regulates the availability of acyl-carnitines in SkM

Given its pleiotropic effects on SkM growth and metabolism, the lower IGF-1 levels in the whole-body PanK4 KO mice confound questions related to the role of PanK4 in skeletal muscle. We therefore generated skeletal muscle-specific PanK4 knockout mice using the Cre-loxP system with skeletal muscle-specific Cre recombinase expression, controlled by the human α-skeletal actin promoter: *Pank4^flox/flox^:HSA-Cre^-/+^* (PanK4 mKO) (Fig 2a). PanK4 mKO mice had normal body weight and food intake (Fig S3a,b), but had twice the amount of fat mass compared to control *PanK4^flox/flox^:HSA-Cre^−/−^* (PanK4 WT) mice (Fig 2b). This was associated with 46% and 70% larger epididymal and inguinal white adipose tissue depots, respectively (Fig S3c). Conversely, PanK4 mKO mice had approximately two grams less fat free mass (Fig 2b); however, this was not accompanied by detectable changes in heart, skeletal muscle, BAT, and liver mass (Fig S3c).

**Figure 2.**
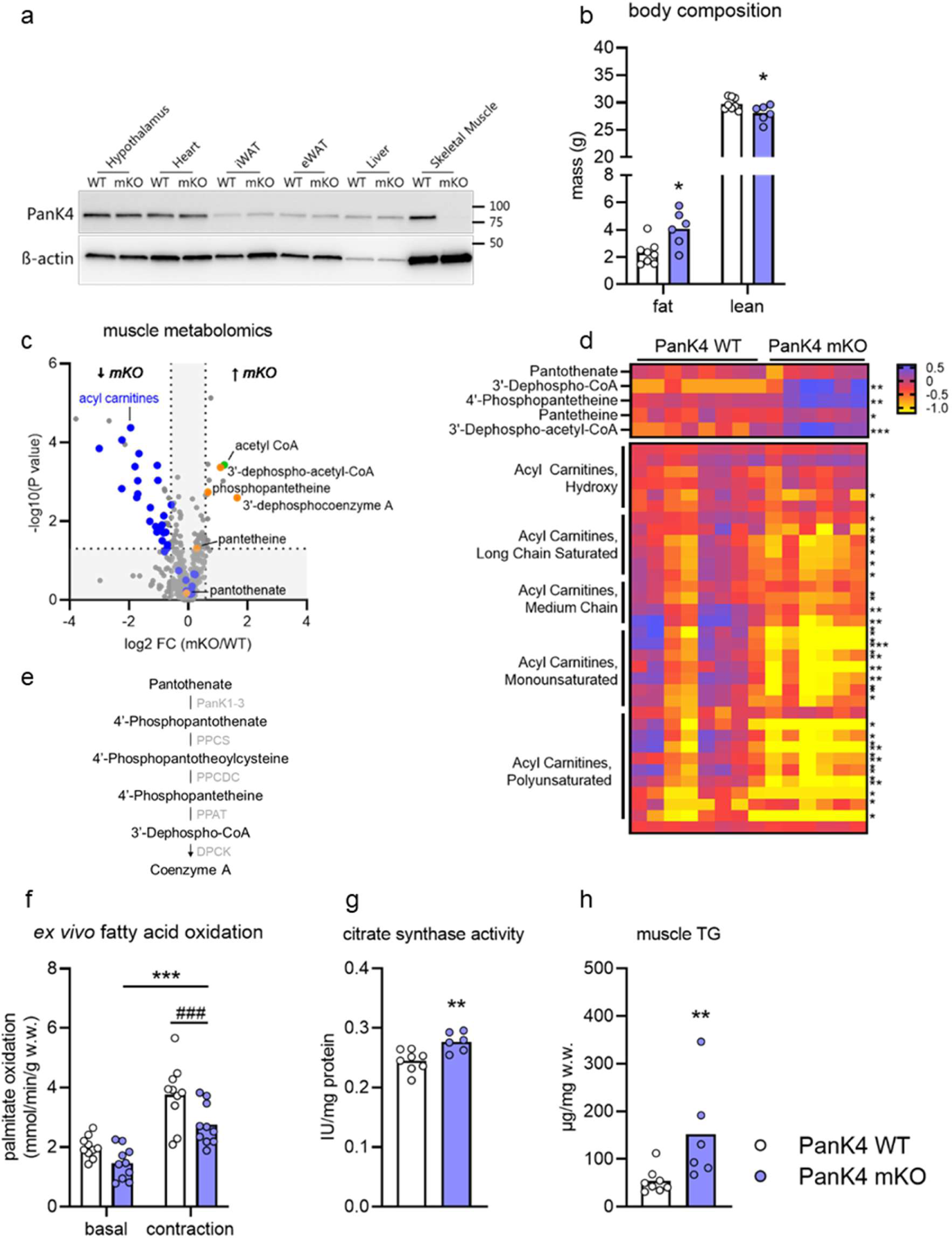
Skeletal Muscle PanK4 regulates fatty acid oxidation. **a,** WB for PanK4 in indicated tissues from male PanK4 wildtype (WT) and muscle-specific PanK4 knockout (mKO) mice at age of 28 weeks. **b,** body composition at age 26 weeks. **c,d,** volcano plot of all 508 detected metabolites and heatmap of indicated metabolites in tibialis anterior muscles from male PanK4 WT and PanK4 mKO mice injected with glucose 40 min prior to euthanizing them at age 28 weeks. **e**, overview of the Coenzyme A biosynthesis pathway. **f,** palmitate oxidation in basal or contracted soleus *ex vivo* from male PanK4 WT and PanK4 mKO mice at age 12-19 weeks. **g,h,** citrate synthase activity in soleus and muscle triglyceride (TG) concentration in gastrocnemius from male mice PanK4 WT and PanK4 mKO mice at age 28 weeks. ***p<0.001, **p<0.01, *p<0.05 vs corresponding PanK4 WT. ###p<0.001 main effect of contracted over basal. Statistic: b, two-tailed unpaired t-test within organ/tissue; c,d, welch test; f, two-way (genotype x contraction) ANOVA; g,h, two-tailed unpaired t-test. Statistical analyses were conducted with log2-transformed data. Šidák post hoc testing was performed whenever respective ANOVA yielded significance. PanK1-3 = pantothenate kinase 1-3; PPCS = phosphopantothenate-cysteine ligase; PPCDC = phosphopantothenoylcysteine decarboxylase; PPAT = phosphopantetheine adenylyltransferase; DPCK = 3’-Dephospho-CoA kinase.

To gain insights into the metabolic function of skeletal muscle PanK4, we performed nontargeted metabolomics in gastrocnemius and tibialis anterior (TA) muscles. Several intermediate metabolites of the CoA synthesis pathway (e.g., 4’-Phosphopantetheine and 3’-Dephospho-CoA) were increased in PanK4 mKO muscles (Fig 2c,d & Fig S3d). This supports recent *in vitro* findings in cancer cells that PanK4 acts as a repressor of the CoA pathway^17^. Strikingly, acyl-carnitines of all types were lower in muscles from PanK4 mKO mice (Fig 2c,d & Fig S3d). Out of the 34 acyl-carnitine species detected, 24 were significantly lower in TA muscles from PanK4 mKO mice than in PanK4 WT, with the remaining 10 being unchanged (Fig 2c,d). Similarly, in gastrocnemius muscle 20 acyl-carnitines were lower and the remaining 14 unchanged in PanK4 mKO relative to PanK4 WT mice (Fig S3d). Levels of both free carnitine and FAs (i.e., acyl groups) were largely unchanged (Fig S3e-g) and circulating FAs were also normal (Fig S3h), indicating that a limitation in precursor supply does likely not account for this striking effect on acyl-carnitines.

### PanK4 regulates fatty acid oxidation in SkM

We hypothesized that lower levels of acyl-carnitine reflects a deficiency in fatty acid entry into the mitochondria, resulting in a decrease in beta-oxidation. To investigate this, we examined fatty acid oxidation (FAOX) using radiolabeled palmitate and the ex vivo isolated incubated muscle technique, specifically focusing on the soleus muscle, because of its higher capacity for FAOX^21^. We investigated both basal- and contraction-stimulated FAOX, because the latter serves as a potent physiological stimulus to increase FAOX. As expected, contraction-induced FAOX was 2-fold higher than the basal rate; however, both basal- and contraction-induced FAOX were 25% decreased in PanK4 mKO muscles (Fig 2f). We next asked whether a decrease in mitochondrial content explained this defect in FAOX in PanK4 mKO mice. Mitochondrial content correlates with the activity of citrate synthase in muscle cells^22^. In soleus from PanK4 mKO, citrate synthase activity was moderately increased (Fig 2g), suggesting that mitochondrial content is unlikely to explain the defects in FAOX. This is supported by similar protein abundance of the different electron transport chain complexes (Fig S3i). Moreover, mRNA levels of proteins involved in fatty acid entry into muscle cells (Fabp3, Fabp4, Cd36, and Fatp4), as well as those related to the carnitine shuttle (Cpt1b and Cpt2) facilitating the mitochondrial entry of fatty acids, were unaltered in PanK4 mKO muscles (Fig S3j). It remains possible that the activity of these proteins is altered. Notably, the reduction in FAOX coincided with an increase in diacylglycerol (DAG) species (Fig 2d and Fig S3k), along with a three-fold elevation in intramuscular triacylglycerol (TG) concentration (Fig 2h). This indicates that instead of undergoing oxidation, fatty acids are shunted towards storage pathways. Collectively, these data indicate that PanK4 appears to play a crucial role in fatty acid metabolism and mitochondrial function, orchestrating efficient fatty acid utilization, with implications for energy metabolism and lipid storage.

### PanK4 regulates acetyl-CoA and metabolic flexibility in SkM

To investigate how PanK4 regulates lipid metabolism, we revisited the metabolomics data. Acetyl-CoA levels were increased by 100% in both gastrocnemius and TA muscles in PanK4 mKO mice (Fig 3a). We were intrigued by this finding because acetyl-CoA levels serve as a critical gauge of cellular fuel, regulating the flux of glucose and fatty acids^23^. We confirmed elevated acetyl-CoA levels in PanK4 mKO gastrocnemius both during fasting and refeeding by targeted analysis (Fig 3b and Fig S4a,b). This was associated with an increase in global protein acetylation in SkM from PanK4 mKO (Fig 3c), consistent with the notion that elevated acetyl-CoA is sufficient to affect the acetylproteome^24^. Notably, skeletal muscle CoA levels remained unchanged in PanK4 mKO muscle (Fig S4c), suggesting that despite the potential role of PanK4 as a repressor of CoA biosynthesis^17^, CoA homeostasis is maintained in skeletal muscle when PanK4 is absent. The expression of mRNAs of PanK1-3, which could act to compensate CoA levels, were unaltered in PanK4 mKO muscles (Fig S4d).

**Figure 3.**
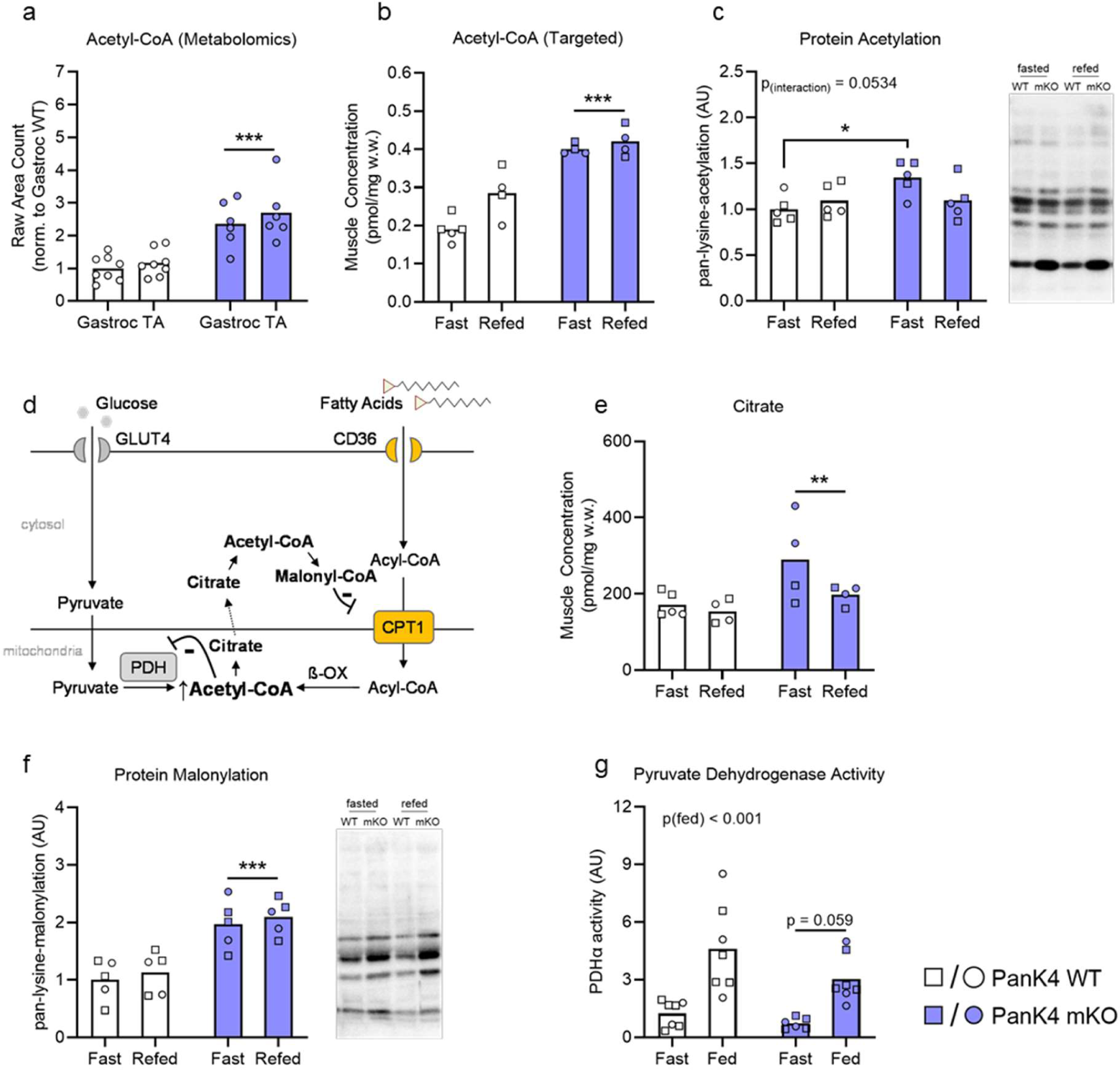
PanK4 regulates acetyl-CoA and metabolic flexibility in skeletal muscle. **a,** acetyl-CoA determined by non-targeted metabolomics in male PanK4 WT and PanK4 mKO mice. **b,** targeted analysis of acetyl-CoA in gastrocnemius from fasted and refed male (circles) and female (squares) PanK4 WT and PanK4 mKO mice. **c**, representative western blots and quantification of pan-lysine acetylation in gastrocnemius muscle from male (circles) and female (squares) PanK4 WT and PanK4 mKO mice. **d**, schematic of how elevated acetyl-CoA could impact metabolism of glucose and fatty acids. **e,** targeted analysis of citrate in gastrocnemius from fasted and refed male (circles) and female (squares) PanK4 WT and PanK4 mKO mice. **f**, representative western blots and quantification of pan-lysine malonylation in gastrocnemius muscle from male (circles) and female (squares) PanK4 WT and PanK4 mKO mice. **g,** pyruvate dehydrogenase activity in gastrocnemium muscle from male (circles) and female (squares) PanK4 WT and PanK4 mKO mice. ***p<0.001, **p<0.01 are main effects; *p<0.05 as indicated (c). Statistic: a,c-e-g, two way (genotype x fast/refed) ANOVA; g, two-tailed unpaired t-test within each indicated gene. Statistical analyses were conducted with log2-transformed data. Šidák post hoc testing was performed whenever respective ANOVA yielded significance.

Persistently elevated mitochondrial acetyl-CoA levels can negatively impact cellular energy metabolism, by leading to the accumulation of citrate, which can be exported from the mitochondria to the cytosol. In the cytosol, it leads to increased levels of acetyl-CoA and malonyl-CoA (Fig 3d). The latter acts as a potent inhibitor of CPT1, a key regulator of mitochondrial fatty acid import. Supporting this idea, our analysis revealed elevated levels of citrate and increased global protein malonylation in PanK4 mKO SkM (Fig 3e,f).

Typically acetyl-CoA levels are thought to be more important for the fine tuning of fatty acid utilization, however, they have also been implicated in the regulation of glucose utilization by allosterically inhibiting pyruvate dehydrogenase (PDH)^25^, an enzyme that facilitates the conversion of pyruvate to acetyl-CoA (Fig 3d). This link might be of particular importance under condition of persistently elevated acetyl-CoA levels as is the case in SkM from PanK4 mKO mice. In accordance with a negative feedback loop being activated, we observed a strong trend (p = 0.059) towards reduced PDH activity in PanK4 mKO skeletal muscle (Fig 3g). These findings collectively indicate that high levels of acetyl-CoA disrupt the efficient utilization of fatty acids, consistent with the observed decrease in FAOX, while also potentially impairing glucose utilization.

### PanK4 regulates fatty acid and glucose utilization in SkM to maintain whole-body insulin sensitivity

Therefore, we also evaluated glycemic outcomes in PanK4 mKO mice. Glucose tolerance was impaired and fasting insulin levels were elevated in PanK4 mKO mice (Fig 4a-c), indicating that skeletal muscle PanK4 is critical for whole-body insulin action. Insulin-stimulated glucose uptake was similar in soleus muscles (Fig S5a) but impaired in PanK4 mKO extensor digitorum longus (EDL) muscles *ex vivo* (Fig 4d). This impairment is not explained by alterations in proximal insulin signaling (Fig S5b-h). We next tested if increasing PanK4 protein is sufficient to boost glucose uptake into skeletal muscle. Using recombinant adeno-associated virus serotype 6 (rAAV6) administered via intramuscular injections (Fig 4f), we overexpressed PanK4 in the glycolytic TA muscle in mice. We first performed a dose-response experiment, assessing the relationship between virus titer and PanK4 protein abundance (Fig S5i) and selected a titer that resulted in a ∼3.5-fold increase in PanK4 protein abundance. Overexpression of PanK4 had no effect on muscle weight (Fig S5j,k). Remarkably, PanK4 overexpression increased basal glucose uptake into resting skeletal muscle by 80% compared to contralateral skeletal muscle (Fig 4g). Skeletal muscle acetyl-CoA decreased when Pank4 was overexpressed (Fig 4h, Fig S5l-n). Collectively, these data suggest skeletal muscle PanK4 is crucial for insulin action and glucose uptake, making it a promising therapeutic target for improving glucose metabolism and combating insulin resistance.

**Figure 4.**
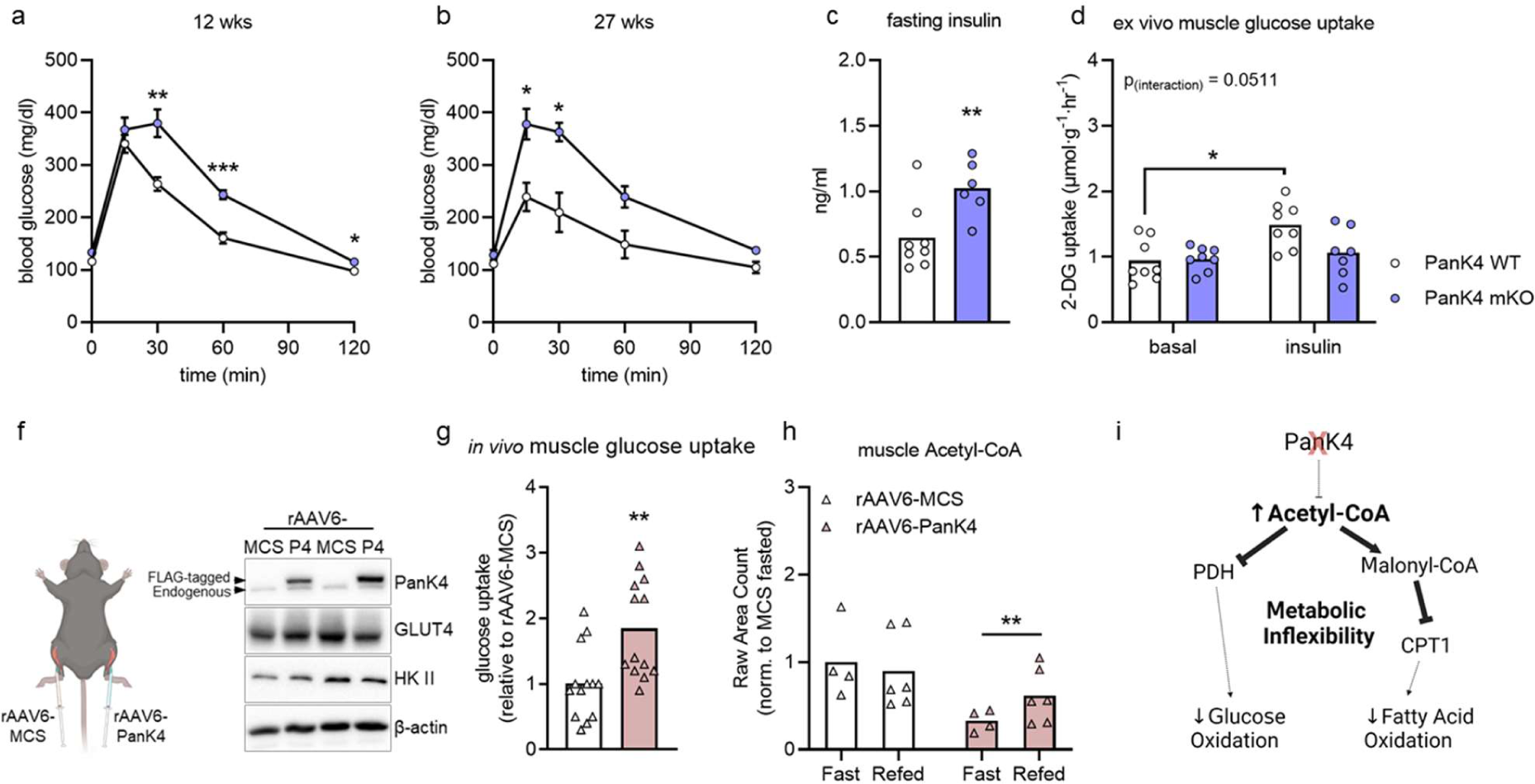
Skeletal muscle PanK4 regulates whole-body insulin sensitivity and muscle glucose uptake. **a,b,** glucose tolerance in male PanK4 WT and PanK4 mKO mice at age 12 and 27 weeks. **c,** fasting plasma insulin concentrations in male PanK4 WT and PanK4 mKO mice at age 12 and 27 weeks. **d,** Insulin-stimulated glucose uptake in skeletal muscle (EDL) from male PanK4 WT and PanK4 mKO at age 13-17 weeks incubated ex vivo under basal conditions or stimulated with 3.0 nM insulin. **f-h,** male C57BL/6J mice age 12-16 weeks were treated with recombinant adeno-associated virus serotype 6 encoding PanK4 (rAAV6:PanK4) injected into tibialis anterior (TA), while contralateral TA was injected with rAAV6:MCS as a control. Virus titer was 1.0E+9 vg. **g, h,** PanK4 protein abundance (f) and glucose uptake (g) in TA muscles were determined 14 days later. **h,** acetyl-CoA determined by non-targeted metabolomics in fasted or refed male C57BL6/J mice overexpressing PanK4 in one TA and MCS in contralateral TA at age 16 weeks. I, schematic for how PanK4 regulates metabolic flexibility via acetyl-CoA ***p<0.001, **p<0.01, *p<0.05 as indicated or vs. corresponding PanK4 WT. Statistic: a,b, repeated measures two-way (time x genotype) ANOVA; c, two-tailed student’s t-test; d two-way (genotype x insulin) ANOVA; g, two-tailed student’s t-test; h, two-way (genotype x feeding) ANOVA. Statistical analyses were conducted with log2-transformed data. Šidák post hoc testing was performed whenever respective ANOVA yielded significance.

## Discussion

We herein identify PanK4 as a highly abundant exercise target in skeletal muscle (SkM) that is involved in the regulation of both glucose and lipid metabolism. Its dysregulation is linked to metabolic disorders and impaired growth and germline deletion of PanK4 impedes normal development. PanK4 regulates SkM acetyl-CoA, facilitating fatty acid oxidation and efficient utilization of glucose. Deletion of PanK4 leads to elevated SkM acetyl-CoA levels, disrupting key metabolic enzymes in SkM and causing whole-body insulin resistance. In a first proof-of-concept experiment, we show that PanK4 is sufficient to enhance glucose utilization in SkM. These findings highlight PanK4 as a novel regulator of SkM energy substrate metabolism with potential implications for combating metabolic diseases.

We focused on exercise signaling because exercise significantly challenges metabolic flexibility in skeletal muscle (SkM). Our interest lay in identifying exercise signals that occur in mice, rats, and humans, as we believed that such conserved responses would be particularly important for SkM metabolism. We discovered PanK4, a highly conserved protein with homologs found in animals, fungi, and plants^16^, as a target in SkM.

The family of pantothenate kinases are known to phosphorylate pantothenate (vitamin B5) to form 4’-phosphopantothenate, the first and rate-limiting step in the CoA biosynthesis pathway. However, mammalian PanK4 lacks this ability and does not contribute to CoA biosynthesis^16, 26^, rendering it a pseudo-pantothenate kinase. PanK4 is also the only family member containing a C-terminal domain of unknown function 89 (DUF89), which can exist independently or, in the case of PanK4, be fused with other proteins^27–29^. It was reported that DUF89 possesses metal-dependent phosphatase activity, targeting phosphorylated metabolites^30^. Recent studies in cultured cell propose that PanK4 dephosphorylate 4’-phosphopantetheine, an intermediate metabolite in CoA biosynthesis, potentially acting as an antagonist of CoA production^17^. Our data indicate the PanK4 antagonizes the CoA biosynthesis pathway also in skeletal muscle in vivo; since, in the absence of PanK4, the levels of 4’-phosphopantetheine increased. However, we observed no significant changes in SkM CoA levels, suggesting that the homeostasis of CoA is maintained through alternative mechanisms in Skm.

Our discovery of an exercise-responsive phosphorylation site on PanK4 complements the recent identification of an insulin-responsive site (T406) on PanK4^17^. However, the functional implications of these phosphorylation sites on PanK4 remain unclear. It would be intriguing to understand whether they affect the phosphatase activity of PanK4. Given that insulin is known as an anabolic stimulus while exercise is considered a catabolic event, it raises the possibility that these two phosphorylation sites integrate competing cellular inputs, fine-tuning the function of PanK4. Undoubtedly, unraveling the regulation of PanK4 in skeletal muscle and other insulin-responsive organs will be of great interest in future research.

In addition, we were astonished by the impact of PanK4 on muscle acyl-carnitine levels, which exhibited significant reductions upon PanK4 loss. Whether this decrease can be solely attributed to a defect in the carnitine shuttle, which regulates the transport of fatty acids from the cytosol into the mitochondria, remains to be elucidated. From a functional standpoint, it is intriguing to note that the loss of PanK4 leads to a 25% decrease in fatty acid oxidation (FAOX) in skeletal muscle (SkM). By comparison, deletion of AMPKα, a recognized key regulator of FAOX in SkM, results in approximately a 20% reduction in FAOX specifically in the soleus muscle^31^. This places PanK4 among the prominent regulators of FAOX in SkM.

PanK4 stands out as one of the few enzymes that plays a crucial role in both glucose and fatty acid oxidation. Typically, there is a reciprocal relationship between the utilization of glucose and fatty acids^25^, meaning that when one is being utilized, it comes at the expense of the other and vice versa. Initially, it seemed challenging to reconcile the fact that PanK4 is required for the efficient use of both substrates. However, we believe this can be explained by the regulation of acetyl-CoA.

Few metabolites can claim a more central and versatile role in metabolism than acetyl-CoA^23^, which is produced during nutrient catabolism to fuel the tricarboxylic acid cycle (TCA), but also functions as a signaling metabolite^23^ and represent a key cellular gauge that regulates energy substrate flux. We propose the hypothesis that, under normal conditions, transient changes in subcellular acetyl-CoA concentration contribute to the regulation of metabolic flexibility by signaling which substrate should be prioritized for fuel production. In this context, persistently high levels of acetyl-CoA, especially in conditions when energy demands are low, could represent a form of cellular overnutrition that disrupts efficient substrate utilization. Essentially, in the absence of PanK4, the energy gauge represented by acetyl-CoA signals a constant state of "fullness," hindering the efficient use of glucose and fatty acids. We show that this metabolic perturbation may in part be explained by impaired function of PDH and CPT1, two key enzymes required for the oxidation of glucose and fatty acids, respectively.

In addition to its role in SkM, it is important to explore the role of PanK4 in the central nervous system. PanK4 exhibits high expression in the hypothalamus and pituitary gland (Fig 1c). Considering the growth/IGF-1 phenotype of the global PanK4 knockout animals, it is tempting to hypothesize that PanK4 plays a role in regulating the release of growth hormone from the pituitary gland. Further studies in this area could provide valuable insights into the involvement of PanK4 in CNS functions.

Overall, we show that PanK4 is a novel regulator of skeletal muscle metabolism that warrants attention as a potential novel therapeutic target to help in the treatment of certain metabolic disease.

## Methods

### Animals Experiments and Housing Conditions

Animal experiments were performed in accordance with the Animal Protection Law of the European Union, and upon permission by the states of Bavaria, Germany or Brandenburg, Germany, or by the Danish Animal Experimentation Inspectorate. Mostly male mice were used, as they are more prone to exhibit dysregulation in glycemia than female mice. In a few critical experiments especially related to the alteration in acetyl-CoA and associated metabolites female mice were included. Mice were group-housed whenever possible. Unless stated otherwise all experiments were performed at ∼22°C with a 12:12 h light-dark cycle. Mice had *ad libitum* access to water and chow diets (Altromin 1324, Brogaarden, Denmark, Altromin 1314, Altromin GmbH, Lage, Germany, or ssniff V1534-300, ssniff Spezialdiäten GmbH, Soest, Germany).

### Human Exercise Study

The detailed experimental procedure can be found elsewhere^32^. Briefly, six male young volunteers (ages 26-28, with BMIs between 23-26 kg/m^2^) participated in a single vigorous cycle session (77%-88% Watt_max_) until exhaustion, which typically occurred after 8-11 minutes of total exercise time. Biopsies from the vastus lateralis were taken before (PRE), immediately after cessation of exercise (0 h post EX) and 2 and 5 hours post-exercise (2 h and 5 h post EX).

### Retrieving GWAS data for PANK4

The GWAS data for PANK4 was obtained downloading information from the NHGRI-EBI GWAS catalog (https://doi.org/10.1093/nar/gkac1010) using FTP servers. By this also not significant associations were received and filtered for a region 50kb up- and downstream of the PANK4 Gene. Subsequently, the Manhattan plot was generated using R-version 4.2.3, utilizing a modified version 0.18 of qqman. This adapted version allowed color coding for variant consequences and replacement of chromosomes with phenotypes. Variant consequences were obtained using the biomaRt R-package version 2.54.

### Mouse Models

#### Pank4^−/−^ (Pank4 KO) mouse

To generate a Pank4 knockout (KO) mouse embryonic stem cell (mESC) the open reading frame of *Pank4* was replaced with a previously published cassette NLS-lacZ-2A-H2B-Venus-neo cassette^20^. Homologous arms (HA) were generated using primers “Pank4 Ex2 fwd” and “PanK4 Ex2 rev” (5’HA, 1919 bp) and “Pank4 Ex19 fwd” and “Pank4 Ex19 rev” (3’HA 1317b bp). The pBKS-NLS-lacZ-2A-H2B-Venus-neo vector was digested with the enzymes, NotI, SacII, SalI and KpnI. Resulting fragments were purified and used together with both HA in Gibson cloning to generate the "Pank4 KO NLS-lacZ-2A-H2B Venus targeting vector." By annealing two single stranded (ss)DNA oligos, two gRNAs with target sites in Exon2 and Exon 19 of Pank4 were cloned into the BbsI site of the pbs-U6-chimaric RNA vector. Both gRNA vectors were mixed with an expression vector of Cas9-nickase (pCAG-Cas9v2D10A-bpA; both vectors were generous gift from O. Ortiz, Institute of Developmental Genetics, Helmholtz München, Germany) and together with the targeting vector were used for transfection of IDG3.2 F1-Hybrid mES cells ^33^. After selection with 300 µg /ml G418, clones were analyzed for correct integration of targeting vector by PCR genotyping and sequencing (primers #1, 2, 3 and 4). Positive targeted mESs were aggregated with CD1 morulae to obtain mouse chimeras. The offspring of the chimeras were genotyped to test for germline transmission of targeted allele (using primers #5, 6, and 7): 790 bp and 559 bp for the wild type or KI allele, respectively). The floxed Neo cassette was deleted by crossing mice with C57BL/6N-Gt(ROSA)26Sor^tm16(cre)Arte^ mice and progeny was genotyped for absence of the selection cassette (using primers #8, 9 & 10): 317 bp and 459 bp for Neo allele product and delta-Neo allele product, respectively. Mice were backcrossed to C57BL/6J background for at least six generations.

**Table 1.**
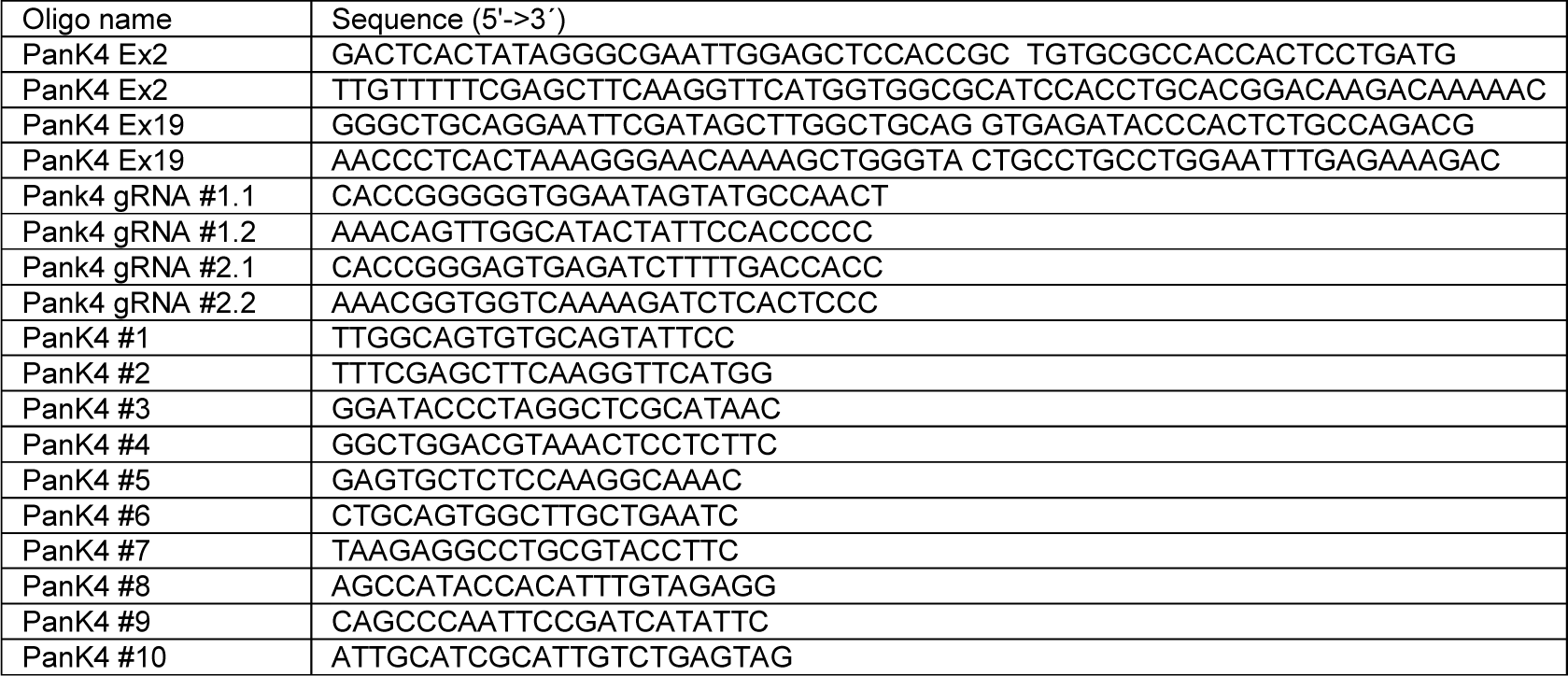
Primers/gRNA used for generating/genotyping PanK4 KO mice.

#### Pank4^flox/flox^ and muscle-specific Pank4 KO (Pank4^flox/flox^:HSA-Cre) mouse

Floxed Pank4 mouse line was created by flanking exon 2 to 9 with loxP sites. Targeting vector constructs were designed as 5’HA(4.3 KB)-loxP-Pank4(exon2-9)-FRT-neomycin-FRT-loxP-3’HA (4.2 KB) using the primers from Table 2. The targeting vector was injected into embryonic stem cells from C57BL/6J background mice, and a Neo-resistant strain was established. Floxed PanK4 mice were obtained from intercrosses with CAG-FLP transgenic mice (C57BL/6N-Gt(ROSA)26Sor^tm1(Flp1)Dym^ . HSA–Cre (stock #006149) transgenic mice were obtained from Jackson Laboratory (Bar Harbor, ME). Homozygous floxed Pank4 mice were then crossed with HSA–Cre mice to obtain muscle-specific Pank4 KO (*Pank4^flox/flox^:HSA-Cre^-/+^*) mice which were verified by PCR, WB and qPCR. Floxed PANK mice only (*PanK4^flox/flox^:HSA-Cre^−/−^*) were used as wild-type (WT) mice in the study. LoxP and HSA-Cre genotyping were done with the specific primers listed in Table 3.

**Table 2.**
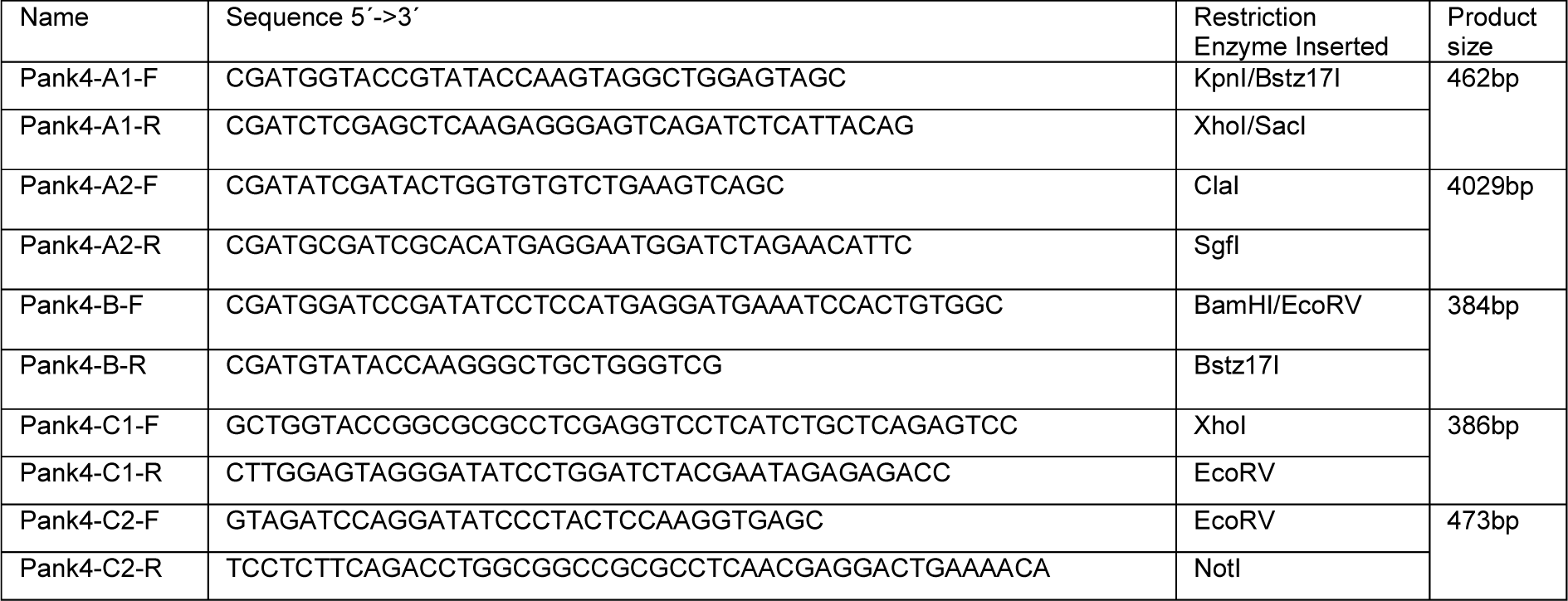
Primers for Floxed PanK4 targeting vector.

**Table 3.**
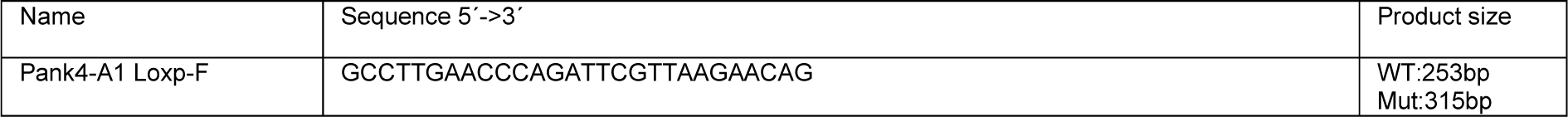
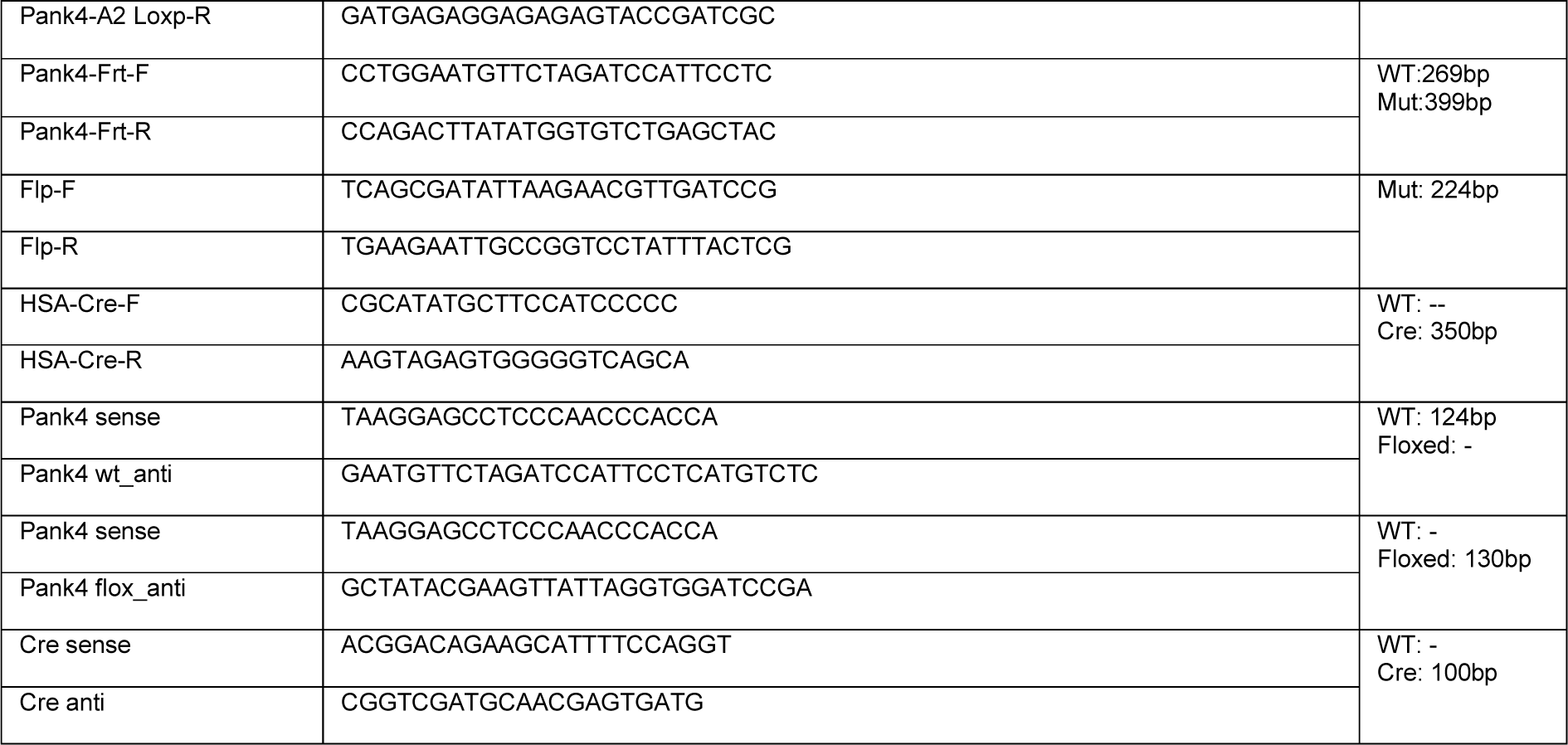
Primers for Floxed PanK4 and HSA-Cre genotyping.

### Gene Expression

RNA from tissues was isolated by Trizol-Chloroform (peqGOLD TriFastTM, 30-2010P, VWR) extraction in homogenized tissues and precipitation with Isopropanol. Genomic DNA was removed (DNAse I, EN0521, Fisher Scientific; RiboLock RNAse Inhibitor, EO0382, Fisher Scientific) and 1µg RNA was transcribed into cDNA using LunaScript RT Super Mix (E3010L, New England Biolabs). Afterwards the mRNA levels were determined by qPCR using SYBR Green (Luna Universal qPCR Master Mix, M3003E, New England Biolabs) on a VIA7 Real Time PCR System (4453534, Thermo Fischer Scientific). Primer efficiency was determined with an experiment-specific standard curve and target gene expression was normalized to mRNA level of housekeeping genes as indicated in the figure legends. List of primers used can be found in Supplemental Table 1.

### Western Blotting

Tissues were homogenized in bead-mill homogenizers in ice-cold homogenization buffer (10% Glycerol, 20 mM Na-pyrophosphate, 150 mM NaCl, 50 mM HEPES (pH 7.5), 1% NP-40, 20 mM β-glycerophosphate, 10 mM NaF, 2 mM PMSF, 1 mM EDTA (pH 8.0), 1 mM EGTA (pH 8.0), 10 μg/ml Aprotinin, 10 μg/ml Leupeptin, 2 mM Na3VO4, 3 mM Benzamidine). Lysate supernatant was obtained by centrifugation for 20 min at 13,000 × g at 4 °C. Lysate protein content was determined with the bicinchoninic acid (BCA) method using BSA standards (Pierce) and BCA assay reagents (Pierce) and all lysates were diluted to the same protein concentration with double deionized water. Total protein and phosphorylation levels of indicated proteins were determined by standard immunoblotting technique loading equal amounts of protein. For determining pan-lysine acetylation a modified buffer was used that contained deacetylase inhibitors: 10% Glycerol, 20 mM Na-pyrophosphate, 150 mM NaCl, 50 mM HEPES (pH 7.5), 1% NP-40, 20 mM β-glycerophosphate, 10 mM NaF, 1 mM EDTA (pH 8.0), 1 mM EGTA (pH 8.0), 10 μg/ml Aprotinin, 10 μg/ml Leupeptin, 2 mM Na3VO4, 3 mM Benzamidine, 1 mM Na-butyrate, and 5 mM Nicotinamide. List of antibodies used can be found in Supplemental Table 2.

### Glucose Tolerance Test

Mice were fasted for six hours from 8 a.m. and intraperitoneally injected with 2 g/kg bodyweight D-glucose (0.2 g in 1 ml saline). Blood glucose concentration in mixed tail blood was measured with a glucometer just before glucose injection (0 min) and 15, 30, 60, and 120 min after glucose injection. At the 0 min time-point extra blood was collected for determination of plasma insulin levels.

### Body composition

Body composition was analyzed using a magnetic resonance whole-body composition analyzer (EchoMRI, Houston, TX).

### Plasma analyses

Insulin was analyzed using the “Mouse Ultrasensitive Insulin ELISA” from Alpco (80-INSMSU-E01). The concentrations of plasma FA (NEFA C kit; Wako Chemicals, Denmark) was measured colorimetrically on an autoanalyzer [Pentra C400 analyzer (Horiba, Japan)].

### Tissue triglycerides

Tissue triglycerides (TG) were determined as previously described^34^ using the kit RandoxTR-210 (Randox) using 20-30 mg of the indicated muscle that was pulverized in liquid nitrogen.

### Fatty acid oxidation

Exogenous palmitate oxidation in isolated soleus muscles of PanK4 and mKO mice was analyzed as previously described^35^. Briefly, the soleus muscles were taken from anesthetized male mice fasted for 3 hours aged 12 – 19 weeks and mounted on a force transducer in a 15 ml container, where they bath at resting tension of 4-5 mN in a Krebs-Henseleit bicarbonate buffer (pH 7.4, 5 mM glucose, 2% FA-free bovine serum albumin, 0.5 mM palmitic acid) at 30°C for 20 minutes. During either rest or contraction (30 Hz, 600 ms pulse duration, 18 tetani/min for 25 minutes), the buffer was exchanged for fresh buffer that also contained palmitate [1-14C] palmitate (0.0044 MBq/ml; Amersham BioSciences, Buckinghamshire, United Kingdom). To seal the incubation chambers, mineral oil (Cat. No. M5904, Sigma–Aldrich) was added on top. At the end of the 25 min long rest or contraction (18 trains/min, 0.6 s pulses, 30 Hz, 60 V) protocol, incubation buffer was collected and the muscles were flash-frozen in liquid nitrogen to determine the rate of palmitate oxidation as described^35^. Palmitate oxidation was determined as CO_2_ production (complete FA oxidation) and acid-soluble metabolites (representing incomplete FA oxidation). As no difference was observed in complete and incomplete FA oxidation between genotypes, palmitate oxidation is presented as a sum of these two forms.

### Citrate Synthase activity

The activity of Citrate Synthase (CS) was analyzed spectrophotometrically as previously described^36^ by tracking the generation of DTNB at 412 nm. The process involved homogenizing soleus muscle in a solution of 50 mM Tris, 1 mM EDTA (pH 7.4) and 0.1% Triton X-100, and then centrifuging the mixture for 10 minutes at 4°C. The supernatant was utilized to measure protein content and levels of CS activity. 10 µl of a 1∶6 diluted tissue sample was placed in one well of a 96-well plate. Afterwards, 215 µl of reaction buffer (100 mM Tris, 1 mM MgCl2, 1 mM EDTA (pH 8.2) and 0.1 M DTNB) and 25 µl of Acetyl CoA (3.6 mM) were added. The procedure was repeated in triplicate. The reaction was initiated by adding 50 µl of Oxaloacetate (3 mM) and the change in absorbance at 412 nm was monitored for 10 minutes at 37°C. The CS activity was calculated based on the slope of the linear part and normalized to mg of protein.

### Pyruvate Dehydrogenase alpha (PDHα) activity

PDHa activity was measured after homogenizing 10–15 mg of wet weight muscle tissue and snap-freezing the homogenate in liquid nitrogen as described previously^37^ and PDHa activity was normalized to creatine concentration in each muscle sample to correct for non-muscle tissue.

### In vitro mouse muscle incubations and insulin-stimulated 2-deoxyglucose (2-DG) uptake

Soleus and EDL muscles from both legs were dissected out from anesthetized mice (6 mg pentobarbital and 0.24 mg lidocaine/100 g body wt). Mice were euthanized by cervical dislocation after muscles had been removed. Muscles were gently lengthened to resting tension (4–5 mN) in incubation chambers (Multi Myograph system; Danish Myo-Technology, Denmark). These chambers contained 4 ml heated (30 °C) Krebs–Ringer–Henseleit (KRH) buffer supplemented with 2 mM pyruvate, and 8 mM mannitol. Muscles were incubated for 30 min without or with insulin (Insuman Rapid, Sanofi Aventis) at a concentration of 1.8 nM and 3 nM for soleus and EDL, respectively. During the last 10 min of insulin stimulation 2-DG uptake was measured with 3H-2-DG and 14C-Mannitol radioactive tracers and 1 mM of 2-DG. Muscles were washed in ice-cold KRH buffer, blotted dry and snap-frozen in liquid nitrogen, trimmed and weighed, before stored at −80 °C.

### rAAV6-mediated overexpression of PanK4 in skeletal muscle

#### rAAV6 generation

Generation of cDNA constructs encoding murine N-terminal Flag-tagged wild-type (WT) PanK4 and a mutant PanK4 in which Ser63 was exchanged with alanine (S63A) were synthesized and subcloned into a recombinant adeno-associated (AAV) expression plasmid consisting of a cytomegalovirus (CMV) promoter/enhancer and SV40 poly-A region flanked by AAV2 terminal repeats (pAAV2) by Genscript (Piscataway, USA) described previously^38, 39^. Co-transfection of HEK293T cells with the above-mentioned pAAV2 and the pDGM6 packaging plasmid generated type-6 pseudotyped viral vectors that were harvested and purified as described ^38, 39^ and are herein referred to as rAAV6:PanK4. In short, HEK293T cells were plated at a density of 8 × 106 cells onto a 15 cm culture dish and 8–16 h later co-transfected with 22.5 μg of a vector genome-containing plasmid and 45 μg of the packaging/helper plasmid pDGM6 using the calcium phosphate precipitate method. Seventy-two hours after transfection, cells and culture medium were collected and homogenized before 0.22 μm clarification (Millipore). The vector was purified from the clarified lysate by affinity chromatography using a HiTrap heparin column (GE Healthcare), ultracentrifuged overnight and resuspended in sterile Ringer’s solution. The puri-fied vector preparations were titred with a customized sequence-specific quantitative PCR-based reaction (Applied Biosystems) (Life Technologies).

### Transfection of mouse muscles using rAAV6 vectors

#### Dose Response

Three months old male and female C57BL/6J mice (Janvier) were placed under general anaesthesia (2% isofluorane in O2) injected into right TA, gastrocnemius and triceps muscle with 1.0 × 10^9^, 2.5 × 10^9^, or 5.0 × 10^9^ vector genomes in a volume of Gelofusine (30 µL) of rAAV6-PanK4 with contralateral muscles receiving corresponding vg of control rAAV6:MCS (multiple cloning site). Mice were euthanized 14 days later and PanK4 and p-PanK4^Ser63^ abundance in indicated muscles were assessed by WB.

#### Glucose Uptake

Three months old male mice C57BL/6J mice (Janvier) were i.m. injected into right TA with 1.0 × 10^9^ vector genomes of rAAV6-PanK4 with contralateral muscles receiving corresponding vector genomes of rAAV6:MCS. 14 days later glucose uptake into TA muscles was assessed *in vivo*. For this, mice were fasted for 2 h, mixed tail blood was collected, before mice were IP injected (10 ml/kg body mass) with saline containing 0.1 mM 2-deoxyglucose (2-DG) and 60 μCi/ml 3H-labeled-2-DG ^40^. Blood was drawn at 20 min post-injection and mice were euthanized by cervical dislocation and TA muscles were rapidly excised and snap-frozen in liquid nitrogen. Plasma 3H activity was determined by scintillation counting and area under the curve from 0 to 20 min was calculated by the trapezoid method to estimate the 3H-2-DG levels that the muscles had been exposed over 20 min [30]. ∼30 milligrams of TA muscle was used to determine the accumulation of phosphorylated 3H-2-DG (3H-2-DG-6-P) with the precipitation method^40^.

### Nontargeted Metabolomics

#### PanK4 mKO samples

tibialis anterior and gastrocnemius muscles from male PanK4 WT and PanK4 mKO mice injected with glucose 40 min pior to euthanizing them at age 28 weeks (see Figure 2).

#### rAAV6-PanK4 overexpression samples

Three months old male mice C57BL/6J mice (Janvier) were i.m. injected into right TA with 1.0 × 10^9^ vector genomes of rAAV6-PanK4 with contralateral muscles receiving corresponding vector genomes of rAAV6:MCS. 14 days later some mice were fasted for 14 h, while others were fasted for 12 h and refed for 2 h by providing free access to chow diet during that time. Mice were euthanized and TA muscles quickly resected out, frozen in liquid nitrogen and stored at −80 degrees Celsius.

#### Muscle Tissue Collection and Preparation for Metabolomics

Muscles were stored at −80 °C and were randomized prior sample extraction. Muscle pieces (∼30–100 mg) were first homogenized with 1.4 mm ceramic beads in water (15 µL/mg tissue) at 10 °C. To extract metabolites and to precipitate the protein, 500 µL methanol extraction solvent containing recovery standard compounds was added to each 100 µL of tissue homogenate. Supernatants were then aliquoted. For *rAAV6-PanK4 overexpression samples*, one aliquot was dedicated for analysis by UPLC-MS/MS in electrospray positive ionization and one for analysis by UPLC-MS/MS in negative ionization. Whereas for *PanK4 mKO samples*, two (i.e., early and late eluting compounds) aliquots were dedicated for analysis by UPLC-MS/MS in electrospray positive ionization and one for analysis by UPLC-MS/MS in negative ionization. Afterward, the extract aliquots were dried under nitrogen stream (TurboVap 96, Zymark) and stored at −80 °C until the UPLC-MS/MS measurements were performed. Three types of quality control samples were included in each plate: samples generated from a pool of human ethylenediamine tetraacetic acid (EDTA) plasma, pooled sample generated from a small portion of each experimental sample served as technical replicate throughout the data set, and extracted water samples served as process blanks.

#### UPLC-MS/MS Non-targeted Measurements

The UPLC-MS/MS platform utilized a Waters Acquity UPLC with Waters UPLC BEH C18-2.1 × 100 mm, 1.7 µm columns, a Thermo Scientific Q Exactive high resolution/accurate mass spectrometer interfaced with a heated electrospray ionization (HESI-II) source, and an Orbitrap mass analyzer operated at 35,000 mass resolution.

#### rAAV6-PanK4 overexpression samples

Two separate C18 columns (2.1 x 100 mm Waters BEH C18 1.7 µm particle) were used: one for acidic (solvent A: 0.1% formic acid in water, solvent B: 0.1% formic acid in methanol) and the other one for basic (A: 6.5 mM ammonium bicarbonate pH 8.0, B: 6.5 mM ammonium bicarbonate in 95% methanol) mobile phase conditions. They were optimized for positive and negative electrospray ionization, respectively. After injection of the sample extracts, the columns were developed in a gradient of 99.5% A to 98% B in 11 min run time at 350 µL/min flow rate.

#### PanK4 mKO sample

For acidic positive electrospray ionization conditions, that chromatographically optimized for more hydrophilic compounds (for early eluting compounds), the extracts were gradient eluted from the C18 column (2.1 x 100 mm Waters BEH C18 1.7 µm particle) using water and methanol containing 0.05% perfluoropentanoic acid (PFPA) and 0.1% formic acid (FA). Another extract aliquot that was also analyzed using acidic positive electrospray ionization conditions, but was chromatographically optimized for more hydrophobic compounds (for later eluting compounds), was gradient eluted from the same C18 column using methanol, acetonitrile, and water; containing 0.05% PFPA and 0.01% FA and was operated at an overall higher organic content. The basic negative ion condition extracts were gradient eluted from a separate C18 column using water and methanol containing 6.5 mM ammonium bicarbonate at pH 8.

The MS analysis alternated between MS and data dependent MS2 scans using dynamic exclusion and a scan range of 80–1000 m/z. Metabolites were identified by automated comparison of the ion features in the experimental samples to a reference library of chemical standard entries that included retention time, molecular weight (m/z), preferred adducts, and in-source fragments as well as associated MS spectra and curation by visual inspection for quality control using proprietary software developed by Metabolon Inc. Only fully annotated metabolites were included for further evaluation. Data were normalized according to raw area counts, and then each metabolite scaled by setting the median equal to 1. Missing data were imputed with the minimum. Biochemicals labelled with an asterisk (*) indicate compounds that have not been officially confirmed based on a standard, but we are confident in its identity.

#### Data Processing and Analyses

Annotations of different experiments have been merged and Welch’s t-test was conducted, comparing two groups to calculate paired or unpaired raw p values. Calculations have been performed using R Version 4. Volcano plots and heatmaps have been created using GraphPad Prism.

### Targeted Metabolite Analysis

Male and female PanK4 WT and PanK4 mKO mice at age 12-17 weeks were overnight fasted for 14 hours. Afterwards some mice had *ad libitum* access to chow diet for two hours while some mice remained without food during that time. Mice were then euthanized and skeletal muscles were quickly resected out.

Approximately 20 mg of mouse gastrocnemius were extracted as follows: 360 µL of 80% methanol containing 0.1 M formic acid and internal standards (listed in Tables S3 and S4) were added to the samples on ice, immediately after addition of extraction solvent the samples were vortexed for 5 seconds and 20 µL of 9% ammonium bicarbonate (w/v in water) were added; the samples were vortexed again for 5 seconds and each sample was shaken with a tungsten carbide bead at 30 Hz for 3 minutes, then the samples were centrifuged at 14,000 g for 10 minutes. The supernatant was transferred to LC vials and dried using a centrifugal concentrator (Thermo Fisher Scientific, Waltham, USA), and finally reconstituted in 50 µL of 50% methanol. Subsequently, the samples were analyzed using liquid chromatography-tandem mass spectrometry (LC-MS/MS) system consisting of an Agilent 1290 UPLC connected to an Agilent 6490 triple quadrupole (Agilent, CA, USA), metabolites were detected in multiple-reaction monitoring (MRM) mode.

The separation and quantification of phosphate-containing metabolites (Table S4, Fig S4a,b) were achieved by injecting 3 µL of the extracts to a HILIC column (iHILIC-(P) Classic, PEEK, 50 × 2.1 mm, 5 µm, HILICON, Umeå, Sweden), the column and autosampler were maintained at 40 °C and 4 °C, respectively. The mobile phase composed of (A) 10 mM ammonium acetate with 5 µM medronic acid in water of pH 6.8 and (B) 10 mM ammonium acetate in 90% acetonitrile. The mobile phase was delivered at a flow rate of 0.35 mL/min at the following gradient elution: 0.0 min (85% B), 5 min (60% B), 7 min (30% B), 8 min (30% B), 9 min (85% B), 15 min (85%). Analytes were ionized in electrospray source operated in the positive mode. The source and gas parameters were set as follows: ion spray voltage 4.0 kV, gas temperature 200 °C, drying gas flow 11 L/min, nebulizer pressure 30 psi, sheath gas temperature 375 °C, sheath gas flow 12 L/min, fragmentor 380 V. Quantification of AMP, ADP, ATP, Mal-CoA, and Ac-CoA was conducted based on calibration curve using their isotopically labeled analogues as internal standards. External calibration curves were used for quantification of Suc-CoA, CoA, and 3-dp-CoA because of unavailability of their isotopically labeled standards. The calibration for all phosphate-containing metabolites covered linear dynamic range from 5 nM to 50 µM.

Carboxylic acids were separated and quantified by using LC-MS/MS after their derivatization with 3-nitrophenylhydrazine (3-NPH). The samples were derivatized according to a published method ^41^ with modifications, briefly: 20 µL of 120 mM EDC (dissolved in 6% pyridine in 50% methanol) and 20 µL of 200 mM 3-NPH (dissolved in 50% methanol) were consecutively added to 20 µL of an extract. The sample was incubated at room temperature (21 °C) for 60 minutes, afterwards 40 µL of 0.05 mg/mL BHT (dissolved in pure methanol) were added to the samples and vortexed. Eventually, 5 µL of each sample were analyzed on an Acquity UPLC HSS-T3 column (100 × 2.1 mm, 1.8 µm, Waters, MA, USA) by using gradient elution of 0.1% formic acid (v/v) in water as mobile phase A and 0.1% formic acid in acetonitrile as mobile phase B – resulting in separation. The mobile phase was delivered on the column by a flow rate of 0.35 mL/min with the following gradient: 0 min (5% B), 12 min (100% B), 13 min (100% B), 14 min (5% B), 17 min (5% B). Column and autosampler were thermostated at 30 °C and 4 °C, respectively. Analytes were ionized in an electrospray ion source operated in the negative mode. The source and gas parameters were set as follows: ion spray voltage −3.5 kV, gas temperature 150 °C, drying gas flow 11 L/min, nebulizer pressure 30 psi, sheath gas temperature 400 °C, sheath gas flow 12 L/min, fragmentor 380 V. The instrument was operated in dynamic multiple reaction monitoring mode (MRM), and the MRM transitions (Table S2) of derivatized carboxylic acids were optimized by using Agilent MassHunter Optimizer. Malate, citrate, pyruvate, 2-oxoglutarate, fumarate, succinate, lactate, and 2-hydroxyglutarate were quantified based on internal calibration with their isotopically labeled standards. Analysis of acids lacking their isotopically labeled analogues (oxaloacetate, isocitrate, cis-aconitate, itaconate) was conducted through external calibration curve. The calibration for all carboxylic acids covered linear dynamic range from 80 nM to 60 µM.

### General Statistical Analysis

Data are presented as means ± SEM in some xy-plots, as individual values only or as means plus individual data points in bar graphs. The statistical analyses of data are described in the figure legends. Graphs were prepared and analyzed with GraphPad Prism Version 9.4.1.

### Funding information

TM and OH are supported by Novo Nordisk Foundation grant NNF18CC0034900. S.H.R is funded by Independent Research Fund Denmark (2030-00007A) and the Lundbeck Foundation (R380-2021-1451). TDM received funding from the German Research Foundation (DFG TRR296, TRR152, SFB1123 and GRK 2816/1), the German Center for Diabetes Research and the European Research Council ERC-CoG Trusted no.101044445. A.M.F. was supported by a postdoctoral research grant from the Danish Diabetes Academy and by the Novo Nordisk Foundation (grant# NNF17SA0031406), and directly by the Novo Nordisk Foundation (grant# NNF22OC0074110). LS is support by the Danish Council for Independent Research, Medical Sciences (grant DFF-4004-00233) and the Novo Nordisk Foundation (grant NNF16OC0023418 and NNF18OC0032082. LLVM is supported by Lundbeck Foundation R322-2019-2688. EAR is supported by the Novo Nordisk Foundation (NNF18OC0034072 II), the Independent Research Fund Denmark grant (DFF-7016-00147), and the Lundbeck foundation (R233-2016-3566). MK is supported by the Deutsche Forschungsgemeinschaft (DFG - KL 3285/2-1), the German Center for Diabetes Research (DZD 82DZD03D03 and 82DZD03D1Y), the Novo Nordisk Foundation (NNF19OC0055192) and the Lundbeck foundation (R288-2018-78).

## Acknowledgement

The authors would like to acknowledge the critical input of Jørgen F. P. Wojtaszewski.

## Supplemental Information

**Supplemental Figure 1.**
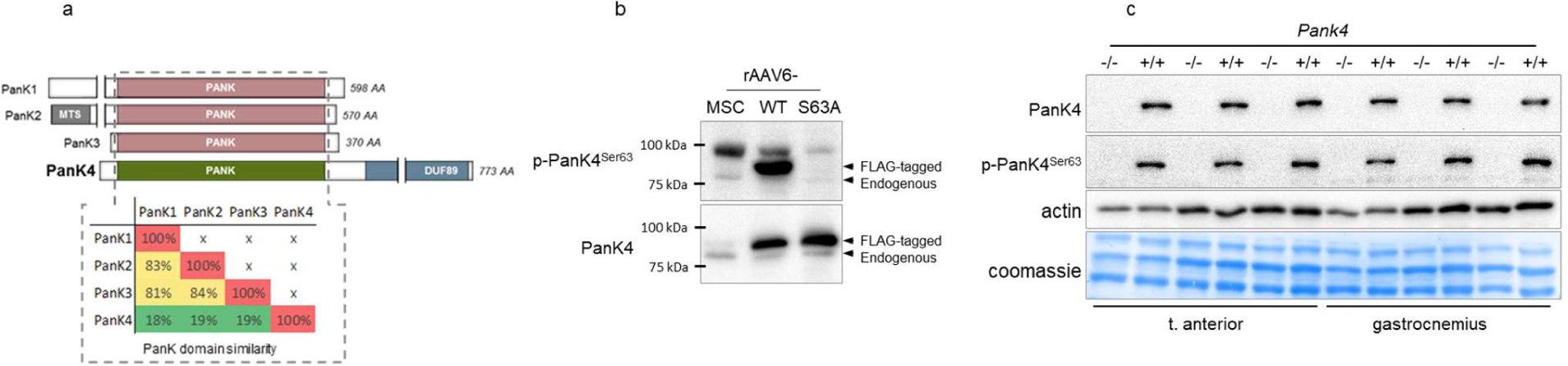
**a,** schematic of pantothenate kinase (PanK) proteins and the %PanK domain similarity at the amino acid level among PanK1-4. **b,** total PanK and PanK4 phosphorylation at Ser63 (p-PanK4^Ser63^) in tibialis anterior muscles from male C57BL/6J mice age 12 weeks that were i.m. injected with recombinant adeno-associated virus serotype 6 (rAAV6) encoding either wildtype PanK4 (rAAV6:WT), a mutant PanK4 in which Ser63 was exchanged with alanine (rAAV6:S63A), or rAAV6:MCS as control. **c,** total PanK and PanK4 phosphorylation at Ser63 (p-PanK4^Ser63^) in indicated muscles from PanK4 WT or PanK4 KO mice.

**Supplemental Figure 2.**
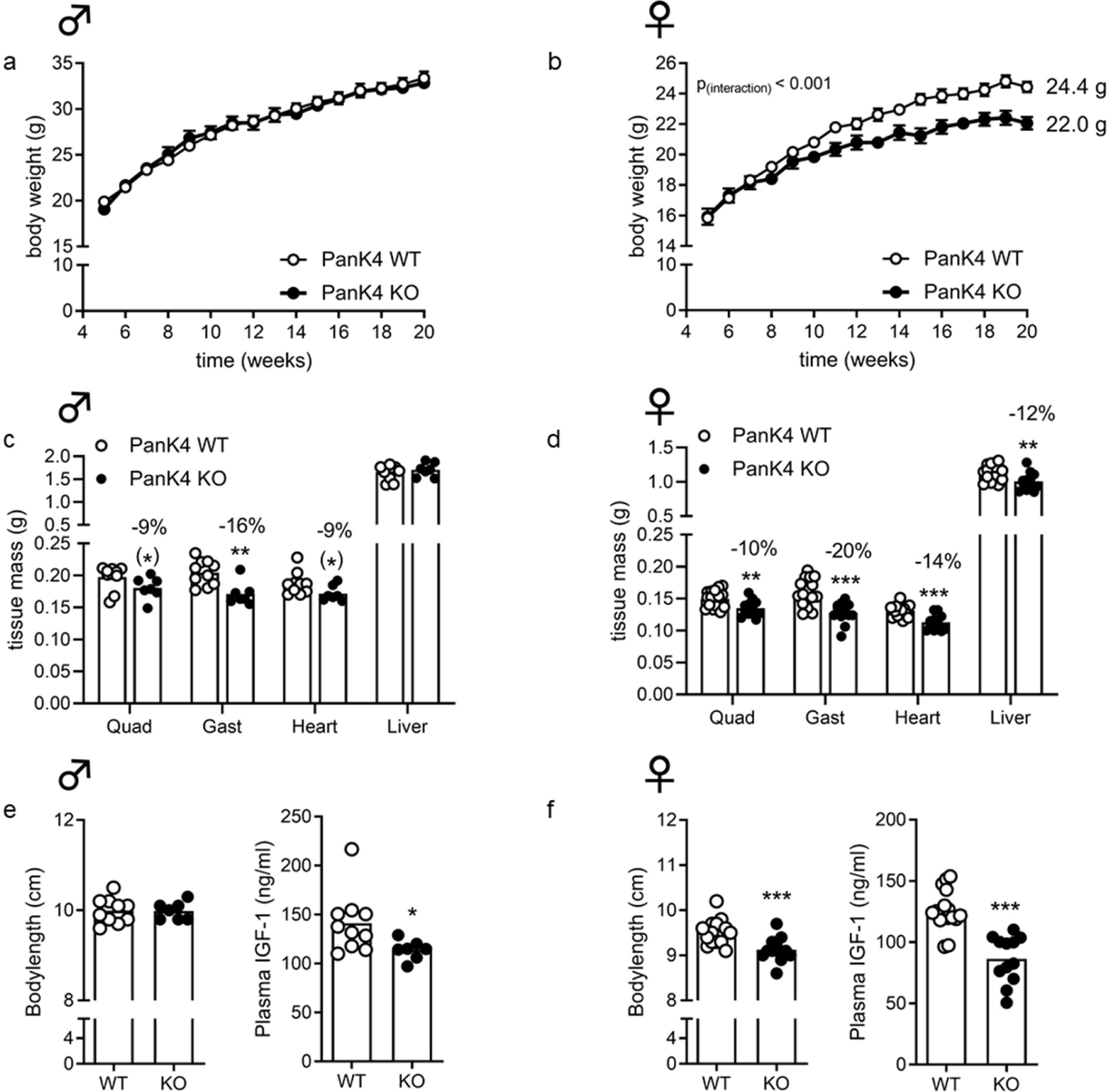
**a,b,** body weight over time in chow-fed male or female PanK4 WT and PanK4 KO mice. **c,d,** mass of indicated tissues in PanK4 WT and KO mice at age 24 weeks. **e,f,** body length and plasma IGF-1 concentration in PanK4 WT and KO mice at age 24 weeks. ***p<0.001, **p<0.01, *p<0.05, (*)p<0.1 vs corresponding WT. Statistic: a,b, repeated measures two-way (time x genotype) ANOVA; c,d, two-tailed unpaired t-tests within tissue; e,f, two-tailed unpaired t-test. Statistical analyses were conducted with log2-transformed data.

**Supplemental Figure 3.**
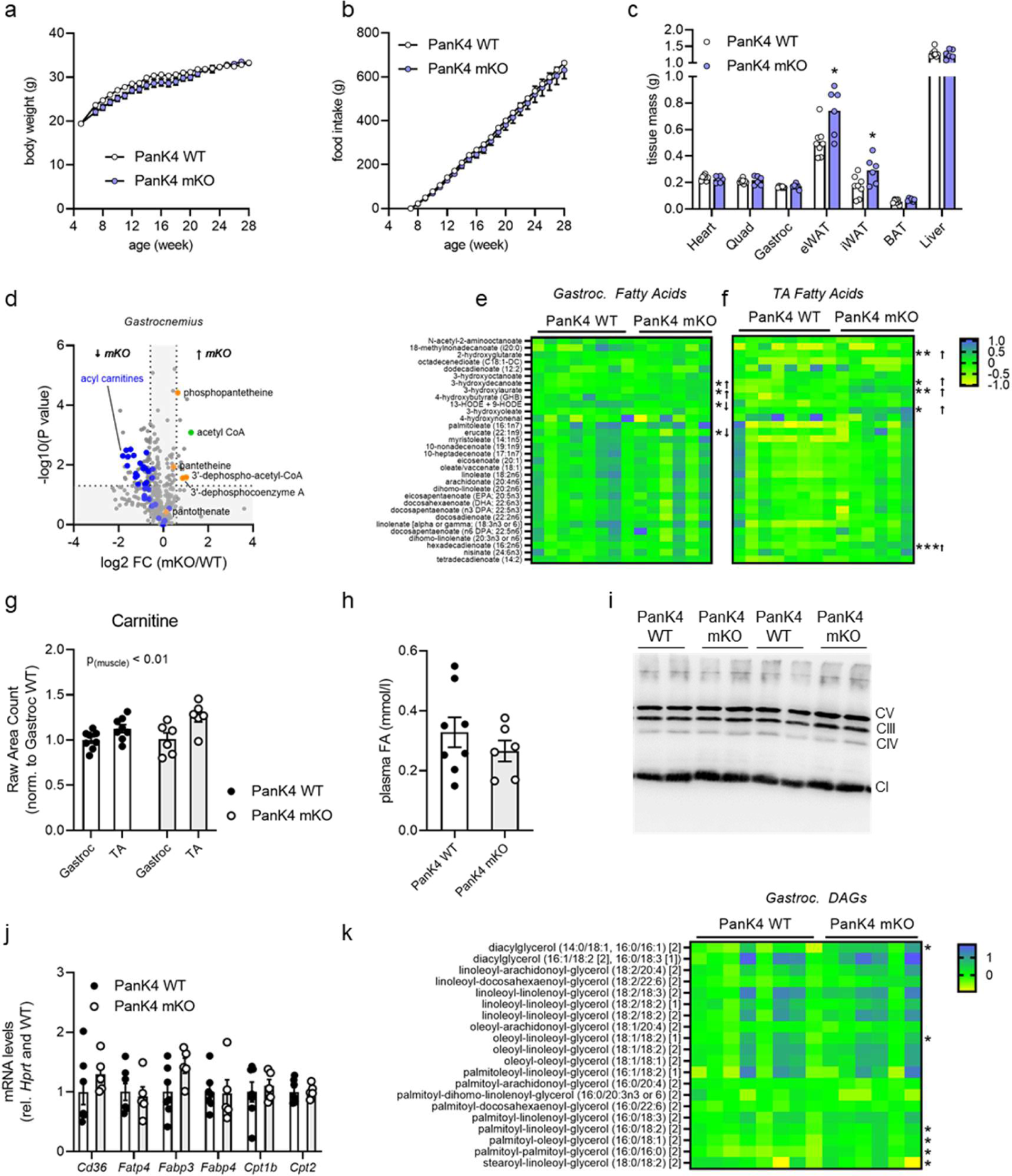
**a,b**, body weight development and food intake of male chow-fed PanK4 WT and mKO mice. **c**, wet weights of indicated tissues from PanK4 WT and PanK4 mKO mice at age 28 weeks. **d-f,** volcano plot (d) of all 508 detected metabolites in gastrocnemius muscles, heatmap (e,f) of indicated metabolites in indicated muscles. **g,** carnitine determined by non-targeted metabolomics in male PanK4 WT and PanK4 mKO mice at age 28 weeks. **h,** plasma fatty acids (FA) from male PanK4 WT and PanK4 mKO mice at age 28 weeks. **i,** OXPHOS western blot in soleus muscles from male PanK4 WT and PanK4 mKO mice at age 28 weeks. **j**, mRNA abundance of indicated genes in gastrocnemius muscles from male PanK4 WT and PanK4 mKO mice at age 28 weeks. **k,** heatmap of indicated metabolites in indicated muscle. ***p<0.001, **p<0.01, *p<0.05 vs corresponding PanK4 WT. Statistic: c, two-tailed unpaired t-test within organ/tissue d-f,k, welch test; g, two-way (muscle x genotype) ANOVA. For c,g, statistical analyses were conducted with log2-transformed data. Šidák post hoc testing was performed whenever respective ANOVA yielded significance.

**Supplemental Figure 4.**
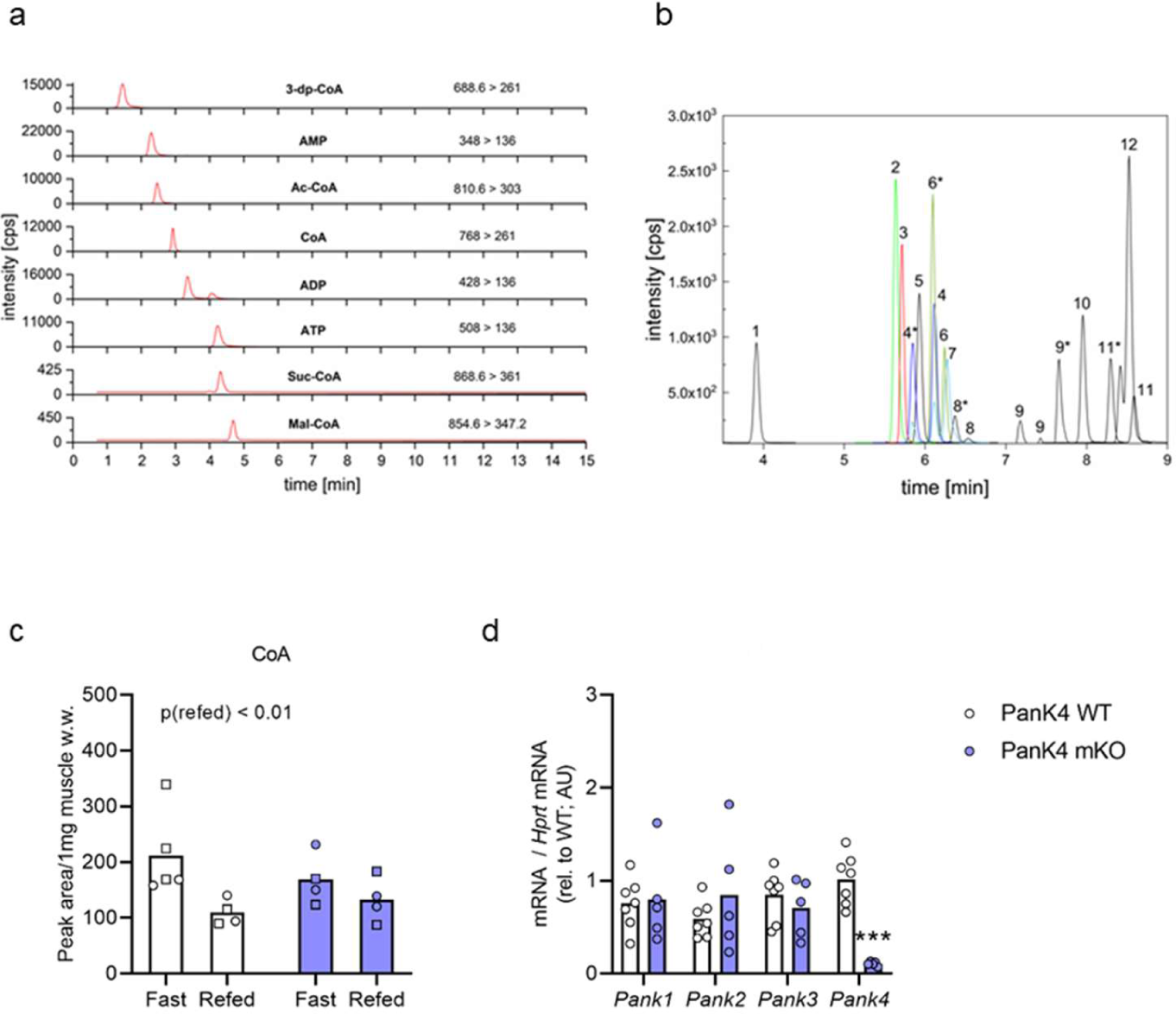
**a,** MRM chromatograms of a 1 µM standard mixture of phosphate-containing metabolites analyzed on an iHILIC-(P) Classic column, 50 × 2.1 mm, 5 µm; **b,** dynamic MRM chromatogram of a 100 nM standard mixture of 3-NPH-derivatized carboxylic acids analyzed on an HSS T3 column. Peak assignment: 1 – lactic acid, 2 – malic acid, 3 – 2-hydroxyglutaric acid, 4 – isocitric acid, 5 – succinic acid, 6 – itaconic acid, 7 – citric acid, 8 – fumaric acid, 9 – oxaloacetic acid, 10 – 2-oxoglutaric acid. Instrumental conditions as described in LC-MS conditions. Asterisk denotes an integration peak used for quantification of carboxylates represented by multiple peaks in chromatogram. **c,** targeted analysis of Coenzyme A (CoA) in gastrocnemius from fasted and refed PanK4 WT and PanK4 mKO mice at age 12-17. **d,** mRNA abundance of indicated genes in gastrocnemius muscles of PanK4 WT and PanK4 mKO mice at age 28 weeks. Statistic: c, two way (fast/refed x genotype) ANOVA. Statistical analyses were conducted with log2-transformed data. Šidák post hoc testing was performed whenever respective ANOVA yielded significance.

**Supplemental Figure 5.**
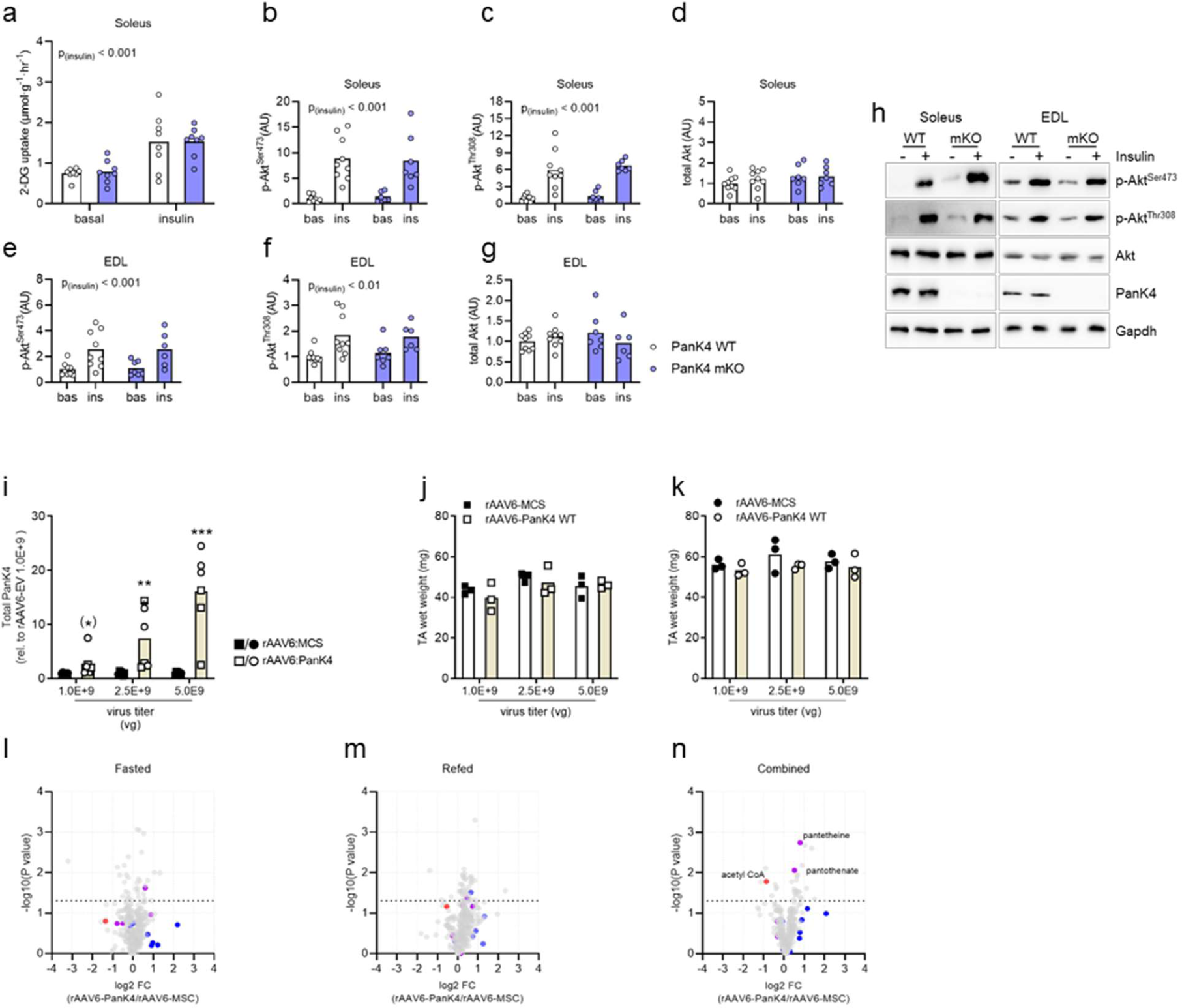
**a,** insulin-stimulated glucose uptake in soleus from male PanK4 WT and PanK4 mKO at age 13-17 weeks incubated ex vivo under basal conditions or stimulated with 1.8 nM insulin. **b-h,** representative western blots and quantification relative to baseline WT of indicated phosphorylation sites or protein in Insulin-stimulated soleus and EDL muscles. **i-j,** dose-response study looking at the effect of virus titer on PanK4 abundance in tibialis anterior (TA) muscle. Male (circles) and female (squares) C57BL/6J mice age 12-16 weeks were treated with indicated titers of recombinant adeno-associated virus serotype 6 encoding PanK4 (rAAV6:PanK4) injected into tibialis anterior (TA), while contralateral TA was injected with rAAV6:MCS as a control. Mice were euthanized 14 days later and (a) PanK4 abundance was assessed by western blot and (b,c) TA mass was determined. **l-n** volcano plot of all detected metabolites determined by non-targeted metabolomics in TA muscles from male C57BL/6J mice age 12-16 weeks treated with rAAV6:PanK4 injected into one TA, while contralateral TA was injected with rAAV6:MCS as a control. Virus titer was 1.0E+9 vg. 14 days later, some mice were fasted for 16 h, while other mice were fasted for 14 h and refed for 2 h. Statistic: a-g, two-way (genotype x insulin) ANOVA with log2-transformed data; i-k, repeated measures two-way (dose x rAAV6) ANOVA with log2-transformed data. Šidák post hoc testing was performed whenever respective ANOVA yielded significance; l-n, welch test.

**Supplemental Table 1.**
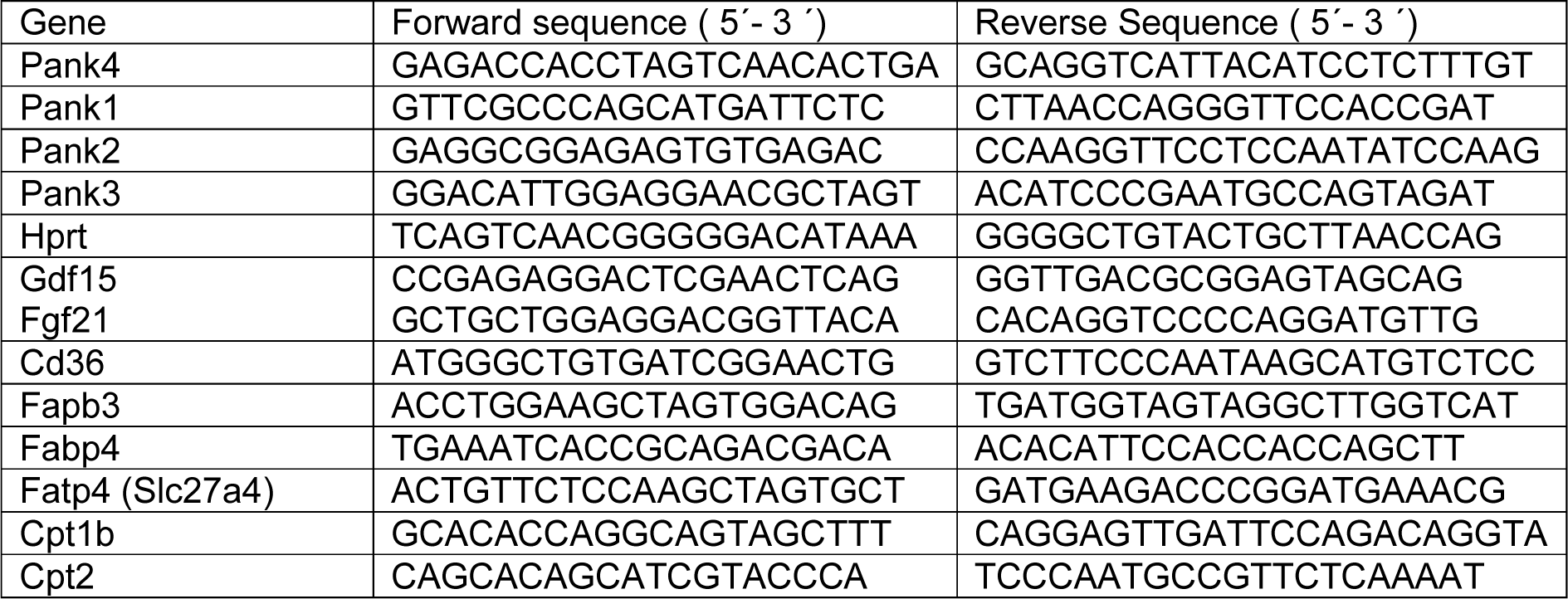

**Supplemental Table 2.**

| Antibody |  |
| --- | --- |
| Pank4 | PANK4 (D8N2C) Rabbit mAb #12055; Cell Signaling |
| Beta-actin | β-Actin Antikörper (C4) HRP sc-47778; Santa Cruz Biotechnology |
| p-PanK4 Ser63 | Custom |
| pan-lysine acetylation | Acetylated-Lysine Mouse mAb (Ac-K-103) #9681s; Cell Signaling |
| pan-lysine malonylation | Malonyl-Lysine [Mal-K] MultiMab Rabbit mAb mix #14942; Cell Signaling |
| p-Akt Ser473 | Phospho-Akt (Ser473) (D9E) XP® Rabbit mAb #4060; Cell Signaling |
| p-Akt Thr308 | Phospho-Akt (Thr308) (244F9) Rabbit mAb #4056; Cell Signaling |
| Akt | Akt Antibody #9272; Cell Signaling |
| Oxphos | Total OXPHOS Rodent WB Antibody Cocktail (ab110413); Abcam |
| GAPDH | GAPDH (G-9) HRP Mouse mAb # sc-365062; Santa Cruz Biotechnology |
| GLUT4 | GLUT4 Polyclonal Antibody #PA-23052; ThermoFisher Scientific |
| Hexokinase II | Hexokinase II (C64G5) Rabbit mAb #2867; Cell Signaling |

**Supplemental Table 3.** Multiple reaction monitoring parameters and retention times of phosphate-containing metabolites.

| Analyte | Retention time [min] | Precursor ion | Product ion | Collision energy [V] | Dwell time [ms] | Type of transition |
| --- | --- | --- | --- | --- | --- | --- |
| **3-dp-CoA** | 1.46 | 688.6 | 261 | 25 | 25 | quantifier |
|  |  |  | 348 | 21 | 25 | qualifier |
| **AMP** | 2.29 | 348 | 136 | 17 | 25 | quantifier |
|  |  |  | 97 | 41 | 25 | qualifier |
| **Ac-CoA** | 2.45 | 810.6 | 303 | 33 | 25 | quantifier |
|  |  |  | 428 | 25 | 25 | qualifier |
| **CoA** | 2.95 | 768 | 261 | 33 | 25 | quantifier |
|  |  |  | 428 | 25 | 25 | qualifier |
| **ADP** | 3.36 | 428 | 136 | 29 | 25 | quantifier |
|  |  |  | 348 | 17 | 25 | qualifier |
| **ATP** | 4.29 | 508 | 136 | 39 | 25 | quantifier |
|  |  |  | 410 | 17 | 25 | qualifier |
| **Suc-CoA** | 4.32 | 868.6 | 361 | 41 | 25 | quantifier |
|  |  |  | 428 | 29 | 25 | qualifier |
| **Mal-CoA** | 4.68 | 854.6 | 347.2 | 34 | 25 | quantifier |
|  |  |  | 303 | 42 | 25 | qualifier |
| **AMP-<sup>13</sup>C<sub>10</sub> <sup>15</sup>N<sub>5</sub>** | 2.29 | 363 | 146 | 17 | 25 | quantifier |
| **Ac-CoA-<sup>13</sup>C<sub>2</sub>** | 2.45 | 812.5 | 305.2 | 38 | 25 | quantifier |
| **ADP-<sup>15</sup>N<sub>5</sub>** | 3.36 | 433 | 141 | 29 | 25 | quantifier |
| **ATP-<sup>13</sup>C<sub>10</sub> <sup>15</sup>N<sub>5</sub>** | 4.29 | 523 | 146 | 39 | 25 | quantifier |
| **Mal-CoA-<sup>13</sup>C<sub>3</sub>** | 4.68 | 857.6 | 350.1 | 34 | 25 | quantifier |

**Supplemental Table 4.** Multiple reaction monitoring parameters and retention times of 3-NPH-derivatized carboxylic acids.

| Analyte | Precursor | Products | Collision energy [V] | Type of transition | Retention time [min] |
| --- | --- | --- | --- | --- | --- |
| **oxaloacetic acid** | 536 | 247.2 | 25 | Quantifier | 8.30 |
|  |  | 357.1 | 25 | qualifier |  |
| **malic acid** | 403 | 208.1 | 17 | Quantifier | 5.64 |
|  |  | 180.0 | 29 | qualifier |  |
| **citric acid** | 443 | 246.1 | 21 | Quantifier | 6.28 |
|  |  | 177.8 | 17 | qualifier |  |
| **isocitric acid** | 443 | 207.9 | 9 | Quantifier | 5.86 |
|  |  | 136.9 | 49 | qualifier |  |
| **pyruvic acid** | 357 | 42.0 | 29 | Quantifier | 7.95 |
|  |  | 136.8 | 29 | qualifier |  |
| **2-oxoglutaric acid** | 550 | 233.1 | 30 | Quantifier | 8.53 |
|  |  | 137.2 | 30 | qualifier |  |
| **fumaric acid** | 385 | 232.0 | 17 | Quantifier | 6.38 |
|  |  | 95.9 | 29 | qualifier |  |
| **cis-aconitic acid** | 578 | 383.1 | 21 | Quantifier | 7.63 |
|  |  | 427.0 | 21 | qualifier |  |
| **succinic acid** | 387 | 234.0 | 21 | Quantifier | 5.93 |
|  |  | 98.0 | 33 | qualifier |  |
| **lactic acid** | 224 | 152.0 | 13 | Quantifier | 3.91 |
|  |  | 137.0 | 21 | qualifier |  |
| **2-hydroxyglutaric acid** | 417 | 137 | 21 | Quantifier | 5.7 |
|  |  | 329 | 21 | qualifier |  |
| **itaconic acid** | 399 | 246 | 17 | Quantifier | 6.1 |
|  |  | 110 | 37 | qualifier |  |
| **2-hydroxyglutaric acid-<sup>13</sup>C<sub>5</sub>** | 422 | 137 | 21 | Quantifier | 5.17 |
| **succinic acid-D<sub>4</sub>** | 391 | 237.0 | 17 | Quantifier | 5.91 |
| **malic acid-<sup>13</sup>C<sub>4</sub>** | 407 | 210.0 | 21 | Quantifier | 5.63 |
| **fumaric acid-<sup>13</sup>C<sub>4</sub>** | 389 | 236.0 | 15 | Quantifier | 6.38 |
| **2-oxoglutaric acid-<sup>13</sup>C<sub>4</sub>** | 554 | 374.1 | 25 | Quantifier | 8.53 |
| **citric acid-D<sub>4</sub>** | 447 | 249.0 | 25 | Quantifier | 6.28 |
| **pyruvic acid-<sup>13</sup>C<sub>3</sub>** | 360.0 | 42.9 | 25 | Quantifier | 7.95 |
| **lactic acid-<sup>13</sup>C<sub>3</sub>** | 227.0 | 151.8 | 13 | Quantifier | 3.91 |

## Notes

https://figshare.com/s/fc5e5d39ac189dbd77f4

### Competing Interest Statement

The authors have declared no competing interest.

